# 3D-printed Planktonic Observational Setup and Analysis Pipeline TrackmateTaxis

**DOI:** 10.1101/2020.08.06.234328

**Authors:** Clemens C. Döring, Harald Hausen

**Affiliations:** Sars International Centre for Marine Molecular Biology, University of Bergen, Norway

**Keywords:** Plankton, Larvae, Behavior, Phototaxis, 3D printing, behavioral assay, analysis pipeline, animal tracking

## Abstract

Planktonic organisms are a cornerstone of marine ecosystems. They vary significantly in size and have a repertoire of behaviors to aid them to survive and navigate their three-dimensional environment. One of the most important cues is light. A variety of setups were used to study the swimming behavior of specific organisms, but broader and comparative investigations need more versatile solutions. With the help of 3D printing, we designed and constructed a modular and flexible behavioral observation setup that enables recordings of animals down to 50μm or up to a few centimeters. A video analysis pipeline using ImageJ and python allows a quick, automated, and robust tracking solution, capable of processing many videos automatically. A modular light path allows the addition of filters or use of pulse width modulation to equalize photon emission of LEDs or additional LEDs to mix different wavelengths. Optionally, a spectrometer can be installed to enable live monitoring of a stimulus. We tested the setup with two phototactic marine planktonic larvae. First, we investigated the spectral sensitivity of the 7-day old larvae of the polychaete *Malacoceros fuliginosus* and second, the behavior of the 200μm spherical bryozoan coronated larvae of *Tricellaria inopinata* to ultraviolet light coming from the bottom of the vessel. The setup and pipeline were able to record and analyze hundreds of animals simultaneously. We present an inexpensive, modular, and flexible setup to study planktonic behavior of a variety of sizes.

## Introduction

Zooplankton is a major factor in carbon and nitrogen cycling within oceans, and plankton migration is influencing movements of other trophic levels in turn (Hays, 2003). Chemical cues, gravity, but most importantly, light are the primary stimuli that aid zooplankton for orientation and navigation within their three-dimensional environment. Light plays an essential role during crucial events of planktonic life stages of numerous animals like dispersal behavior or during the transition from the pelagic larval phase to the adult benthic, often sessile stage.

Additionally, the daily vertical migration of zooplankton is primarily controlled by light (Thorson, 1964). Respective directional movements towards and away from light (phototactic steering) are well documented for many animal larvae like sponge larvae (Elliott et al., 2004), Coral (Piraino et al., 2011), Nauplius (Swift & Forward, 1980), annelid and mollusk trochophore (Jékely et al., 2008; Marsden, 1986; Miller & Hadfield, 1986) but also adult stages like copepods (Tranter et al., 1981), Cladoceri (Ringelberg, 1964) and Daphnia (Effertz & von Elert, 2014). Also, light-dependent swimming behavior can be irrelevant to the direction of the stimulus like the UV avoidance reaction of *Platynereis dumerilii* larvae (Veraszto et al., 2018), *Drosophila melanogaster* (Xiang et al., 2010), *Salmo salar* (Fernö et al., 1995) or the shrimp *Mysis relicta* (Gal et al., 1999). However, for the vast majority of planktonic animals, many essential aspects are poorly understood, such as the specific reactions to different wavelengths, the switching between positive and negative phototaxis, the influence of chemical cues, or other factors on light-dependent reactions and the changes of all of this throughout development. A better characterization of light-dependent movements of planktonic organisms is undoubtedly relevant for a deeper understanding of planktonic ecology. Still, it will also affect research on the behavioral and sensory biology of planktonic organisms. For a few animals, specialized setups and pipelines for assessing and analyzing the behavior have been developed. In this study, we present an affordable, modular setup for quantitative analysis of the swimming behavior of many different kinds of planktonic organisms. The main focus is on light-dependent behavior, but the setup can easily be modified for studies on chemo-sensation or gravitaxis.

Assessing the impact of light on the planktonic movement within the water column consists of two separate challenges. The first is to observe the animals in near-natural conditions while still being able to obtain quantifiable, reproducible data about swimming behavior. The second challenge deals with quantification and processing the generated data.

1. Observation: Planktonic animals are diverse and range from only a few micrometers up to several meters. Our interest lay with the micro-up to mesozooplankton. The setup was supposed to be able to record animals from only 100μm up to a few millimeters. Most published setups for behavioral experiments are custom constructions for a specific organism and would be limited in its flexibility. A huge variety of installations for behavioral observations exists for the most common model organisms, but for marine organisms, the selection is limited. For fish, varying size constructions exist to observe navigation in rivers (Rodríguez et al., 2015) or tracking animals in 3D with the help of sensor bars from Microsoft Kinect (Saberioon & Cisar, 2016). For even smaller marine animals like plankton, some setups are capable of observing animals in situ with the help of a remotely operated vehicle (Gallager et al., 2004) or for simulating the mangrove environment of the boxed jellyfish *Tripedialia cystophora* (Garm et al., 2007; Oskarsson et al., 2015; Petie et al., 2011). Even smaller zooplankton of a few millimeters and smaller setups have been used to analyze the swimming behavior of trochophore larvae of *Platynereis dumerilii* under various light conditions (Veraszto et al., 2018) or to observe escape (Fields & Yen, 1997) or mating (Kiørboe, 2007) behavior of copepods. The often sophisticated setups used in these and other studies are often not described in such detail that they can quickly be rebuilt or are not flexible enough to allow usage with different animals or varying stimuli. Our focus was to design a basic setup that is easy to build, allows precise adjustment of various light conditions, and which can be easily modified and extended by adding further modules, e.g., for chemical stimulation.
2. Quantification: Behavioral assays generate large quantities of data that are not feasible to process manually. For this purpose, an extensive selection of software exists with varying degrees of complexity or specialization. Often, tracking software is specially designed for specific animals like MouseMove (Samson et al., 2015), EthoWatcher (Crispim Junior et al., 2012) for mice, Ctrax (Branson et al., 2009) for Drosophila or a plethora of trackers for *Caenorhabditis elegans* (for an overview see Hart, 2006). Others focus on adding further functionality; in particular, the problem occurring when tracked animals cross paths has been addressed with varying degrees of success (toxtrac (Rodriguez et al., 2017), idtracker (Pérez-Escudero et al., 2014)). While these trackers allow tracking of individual animals, it is still recommended to keep animal numbers low to avoid too many crossing events to happen at the same time.

Aiming higher experimental throughput, we provide a simple, automatic, and robust pipeline to assess directional swimming behavior from recordings of high numbers of animals, applicable to various kinds of swimming trajectories and organism size. All steps run within graphical user interfaces, from video preprocessing to the generation of plots and statistical output. It makes use of the particle tracker Trackmate within Fiji, but tolerates high amounts of path splitting and joining events.

Planktonic behavior within the water column is complex, and different stimuli influence such behavior. We describe an experimental setup to investigate the light-dependent swimming behavior of a wide variety of zooplankton under freely configurable light conditions. By utilizing 3D printing, we demonstrate a straightforward and affordable way to add functionality to this setup for more specific investigations and studies of other stimuli. Furthermore, we supply an easy software pipeline for data analysis.

## Materials and Method

We designed all the parts for 3D printing with FreeCad (https://www.freecadweb.org/) and printed in ‘Black Strong & Flexible’ plastic by shapeways.com. All model files are available in various file formats. For the backlight illumination, we utilized an IR backlight (Polytec, Waldbronn, Germany, SAR7 0605) and its controller (Polytec, Waldbronn, Germany PAD2 2135/1) to adjust illumination intensity finely. The camera used in our setup is a monochromatic CMOS USB3 Camera (Imaging Source, Bremen, Germany, DMK 23UX236) together with a lens (Ricoh, Tokyo, Japan, FL-CC1614-2M). Optionally, we mounted a near-infrared longpass filter (Reichmann Feinoptik, Brokdorf, Germany, RG610, 03.1.1.1.0702) in front of the camera, to avoid light scattering by the illuminating stimuli during phototaxis experiments. To mix different wavelengths of light, we used dichroic filters, beamsplitters, and mirrors from Omega Optical (Brattleboro, USA, 430DCLP, 505DRLP, 565DRLPXR, 50/50BS) in connection with LEDs from Pur-led (Undenheim, Germany, for full details, see “Building Instructions” in supporting information). The LEDs were either powered by a standard laboratory power supply (Tenma, Osaka, Japan, 72-10505) or with an Arduino Uno Rev3 (Arduino.cc) microcontroller and appropriate resistors of the E24 series. To be able to fine-tune light intensity, we utilized neutral density filters from Reichmann Feinoptik or pulse width modulation. We used a spectrometer (B&W TEK, Newark, USA, SpectraRad, BSR112E-VIS/NIR) to monitor the irradiance of the emitted light. Additionally, we also measure LEDs emitted spectrum with Ramses Spectrometer (TriOS, Rastede, Germany, 40S121010). To avoid any light contamination, we encased the whole setup in a simple wooden frame that was covered by a 0.12mm thick blackout fabric (Thorlabs, Newton, USA, BK5). Scripting was done in Fiji version java6 20170530 and python version 2.7 with Anaconda2 version 4.0.0 using spyder2. The script is also available in python 3.8

### Animals

We collected the bryozoan *Tricellaria inopinata* colonies in Brest, France. Aquariums in light-tight boxes housed the adult colonies at a constant 18° while maintaining a strict 12hr day/night cycle. Light induces spawning of larvae. Each experiment started 30minutes after spawning began to ensure a similar age of larvae involved. Larvae of the annelid *Malacoceros fuliginosus* were reared from our lab culture.

## Results

### Overview

3D printing has been an emerging technique used for a variety of custom build applications. It has been used to construct microscopes (Rawat et al., 2017), chemical reaction chambers (Symes et al., 2012) or microfluidic devices (Erkal et al., 2014). Utilizing 3D printing has the advantage that it is available in different types of polymers and other materials, and it is possible to construct complex parts without the use of heavy or expensive machinery or tools. 3D printers are commercially available and more and more widespread. Furthermore, companies offer to print 3D models using a variety of methods and make owning the printer, not a necessity. We ended up using a commercial 3D printing service and opting for selective laser sintering (SLS) printing due to its rougher surface finish and that this method makes the use of support structures during the print (scaffolding) unnecessary.

The behavioral setup consists of a camera mounted on a slider within a track (Fig 1 A). All main parts, except for the cuvettes, can be printed from the 3D models provided in the supporting information. Printing can be performed on locally available 3D printers or by commercial suppliers of 3D printing services. Detailed instructions for the final assembly are provided in the supporting information “Detailed Building Instructions”. Camera height and angle are adjustable by simple thumbscrews. At the end of the track is the fixture to hold a cuvette. The cuvette holder can be mounted in 7 different heights and is capable of holding a cuvette of varying depth up to approximately 18cm high, 23cm wide. Above and below the cuvette holder are guided tracks to hold a light path or other fixtures in place. Positioned behind the cuvette is an IR backlight illuminating the observable area. The camera track can be constructed to be either 23, 46, or 69cm long and allows movement of the camera closer or further away from the observable area depending on cuvette size and animal size.

**Figure 1.**
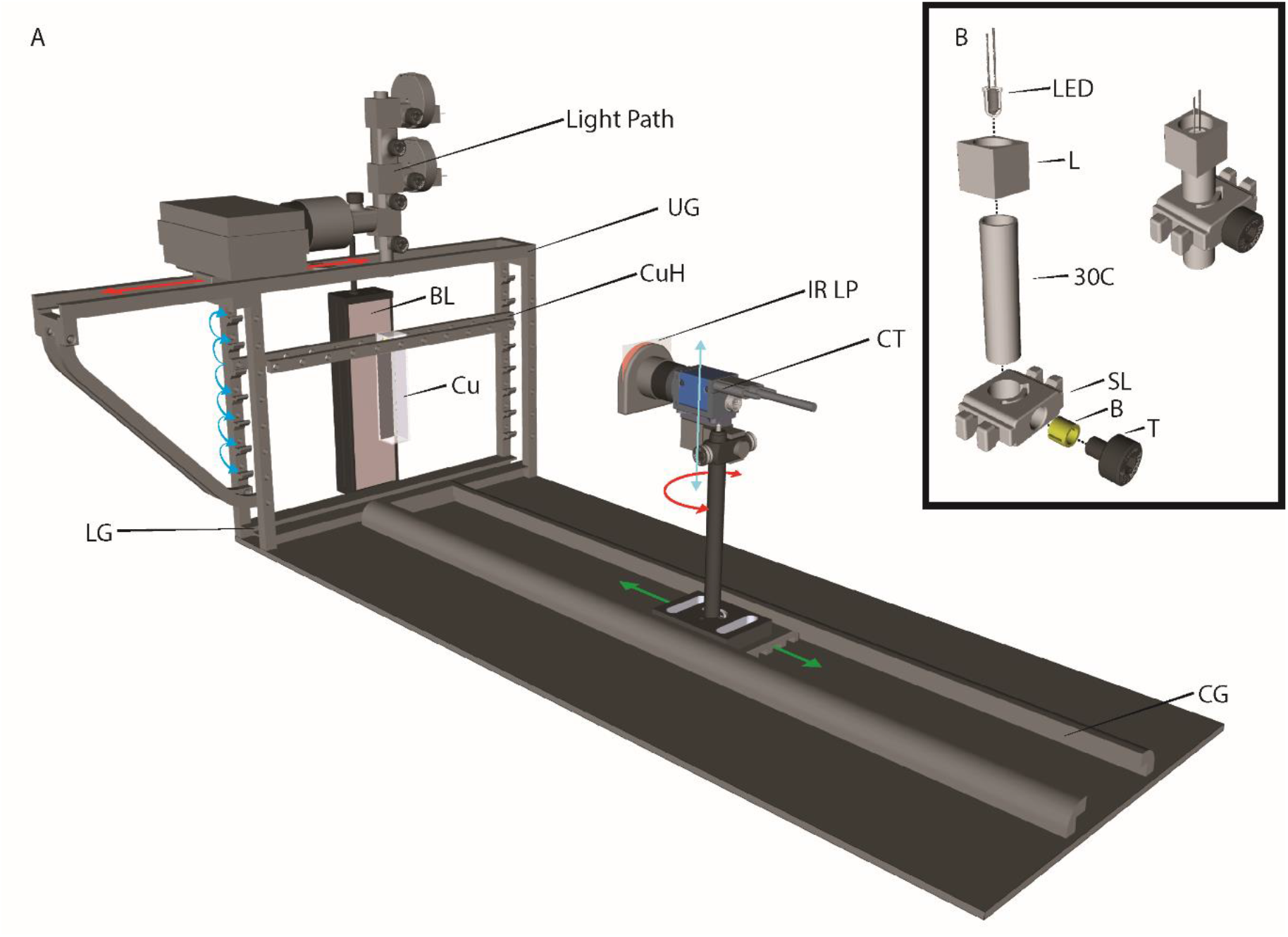
Overview of the Setup. A. The setup consists of a camera with an infrared longpass filter (IR LP) mounted on the camera tower, which can be adjusted in height (blue arrow), rotated (red circle), and variable working distance (green arrow). The camera observes a cuvette (Cu) that can be mounted at varies heights (Blue curved arrows) and receives backlight infrared illumination (BL). As a light stimulus, various light paths can be constructed that are mounted either above or below the cuvette. B. A very simple light path can be built with few pieces that allow an adjustable height of the LED. BL: backlight, Cu: cuvette, CuH: cuvette holder, LG: lower guide, UG: upper guide, CG: camera guide, CT: camera tower, IR LP: infrared longpass filter, L: LED cube, 30C: 30mm connector, SL: slider, B: brass inlet, T: thumbscrew

### About Camera, Animals, and Cuvettes

Essential parameters for any setup depend on the animal to be studied. Most important are appropriate dimensions of the cuvette allowing for near-natural swimming behavior. In turn, camera optics have to match cuvette dimensions in x, y, and z to achieve sharp images from anywhere within the cuvette, and camera-resolution has to be high enough that individual organisms can be tracked. With relatively simple calculations, it is possible to determine the desired characteristics and physical limits of any given camera, associated lens, and cuvette based on the animals’ size.

#### Animal size consideration

When it comes to determining an optimal camera and lens, its magnification factor is, of course, the essential property considering that most planktonic animals are microscopic. The property that concerns us the most is how many pixels our animal will occupy on any given chip.

This height of the object under investigation (h^o^) relates according to basic optic laws to focal length (f), distance to the object (d) and the size of each pixel on the sensor (h^p^):

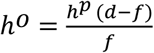

The camera we installed has a CMOS chip of 1/2.8 inches and a pixel size of 2.8 μm. The lens we used has a focal length of 16mm and a minimum working distance of 18 cm. Since our setup positions the camera on a continuous slider, the camera position can vary up to a maximal distance of 60cm. If the camera position is at a minimum working distance to the cuvette, each pixel on the camera will picture 28 μm. In its furthest configuration, each pixel translates to 100 μm. Theoretically, this setup is capable of recording animals anywhere between 28 and 100 μm as its lower limit. However, we do not recommend using a resolution of one pixel per animal, as this can cause a lot of noise later when tracking the animals. Even minute changes of only one pixel will cause the tracker to identify false-positives. Considering that each animal should at least occupy 2×2 pixels, it would limit our camera-setup to animals roughly 50μm in size. This value can easily be adjusted by using cameras with other pixel sizes or different lenses.

#### Depth of field

Another critical parameter of the camera optics for obtaining sharp images is the depth of field (DOF). It relates to working distance (d), the circle of confusion (c), the f-Number (N), and the focal length (f):

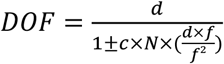

The circle of confusion in the case of a digital camera is the pixel size of the sensor, which is 2.8 μm in our case. As the formula implies, the focal length of the lens influences our depth of field enormously. This should be taken into account when deciding on the lens to be used. For our camera and lens, the depth of field is between 2.5mm and 30mm with an aperture of 4. Further closing the aperture will increase the depth of field slightly but requires more illumination. While these values reflect the theoretical limits of the depth of field, other factors influence the effective, workable volume. Image processing and the capabilities of the tracker can compensate for recordings, which are slightly out of focus. For our setup, it was possible to effectively track animals in a volume up to four times deeper than the calculated depth of field.

#### Observable area

Finally, the visible area changes with the position of the camera and limits the size of any cuvette used. The observable area is directly dependent on the camera chip resolution in pixels(w^i^ × h^i^), the object hight that each pixel is capable of recording (h^p^) and the working distance (d):

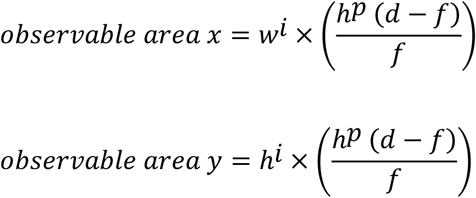

The camera we used has a full HD chip resolution of 1920×1200 pixel. In the closest possible configuration of our setup, the observable area is 5.5cm × 3.4cm, while the furthest possible position is capable of recording an image of 21.6cm × 13.5cm in size.

### Lightpath

#### Light source

Several options exist for generating light of a defined wavelength and intensity. Monochromators are convenient for narrowing down the spectrum of a light source to a narrow band at the desired wavelength. However, they are costly investments and provide only one band at a time. Additionally, the emitted energy (irradiation) may vary across the spectrum depending on the spectral characteristics of the used light source. Commercial LED light for microscopic purposes is an attractive alternative, but they are likewise costly and may not offer the wavelengths desired. Simple LED light bulbs are cheap and available in many colors and intensities. Thus, we decided to use a LED-based light path, which allows for illuminating the samples with light composed of several wavelengths and at desired intensities.

Our setup is modular and allows many different configurations. A very simple light path could use a single LED as a light source. To provide standardized light conditions, this rests inside an LED holding cube connected to a 12mm wide tube that is fixed to a slider either at the bottom or top guide tracks (Fig 1 B). A more complex light path could consist of multiple LEDs that merge with the help of dichroic mirrors (Fig 2, supporting information “Detailed Building Instructions”).

**Figure 2.**
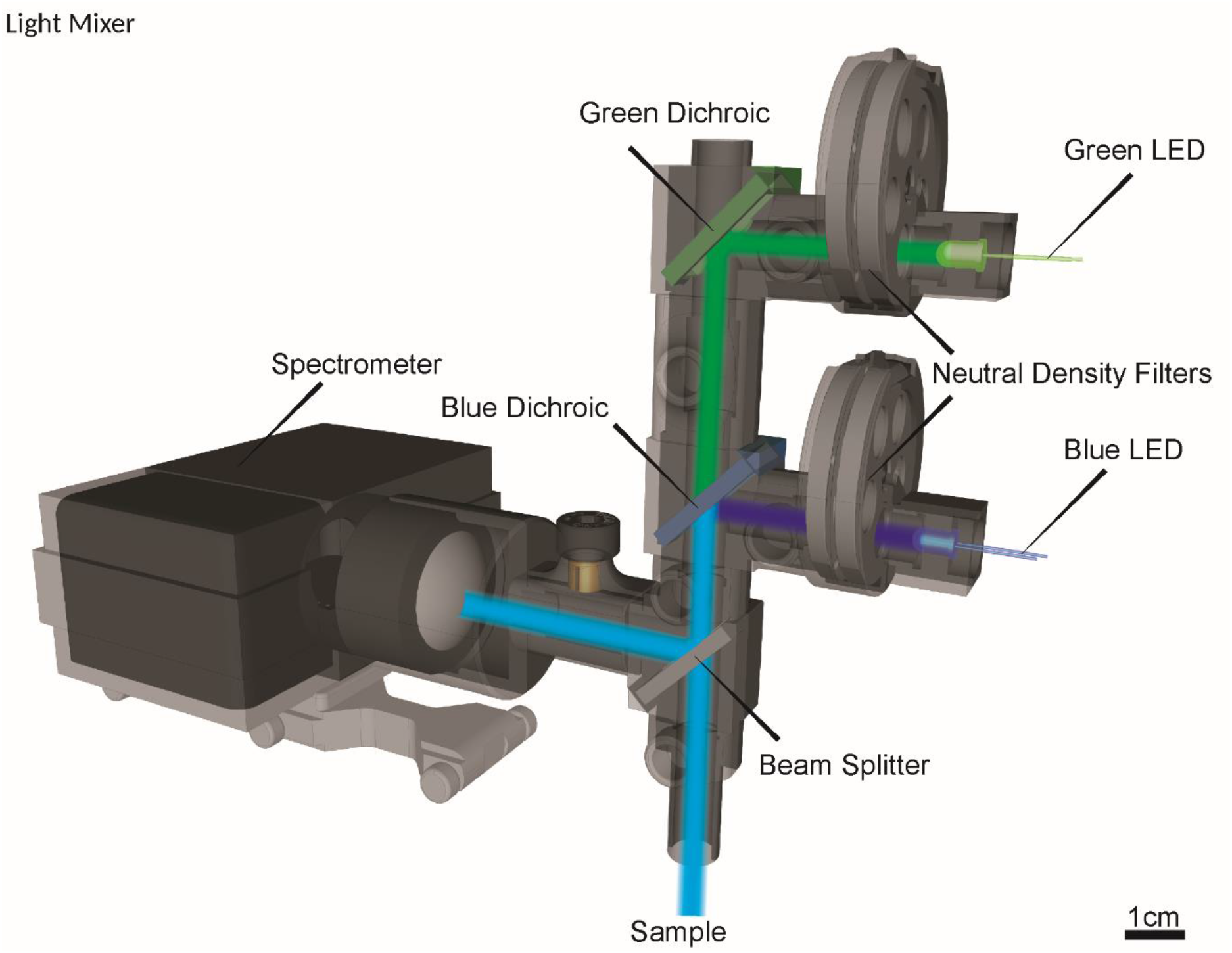
The light mixer allows the mixing of multiple LEDs (shown here with two) that can all be individually regulated with neutral density filters inside of the filter revolver and simultaneously measuring irradiance with a spectrometer.

#### Controlling photon emission

For adjustment of overall light intensity and that of specific wavelengths in mixed light conditions, the output of each LED has to be controlled. A prevalent option to regulate the intensity of LEDs is to flicker them at high frequencies by pulse-width modulation (PWM) of the voltage supplied. PMW is a convenient approach to adjust the average irradiance of LEDs over a specific range, which depends on the electronic characteristics of the controller, the LEDs, and the flicker frequency. When dimming with PWM, it is desired to have a high contrast ratio, which can be achieved with low-frequency PWM. However, the flicker fusion frequency of the eyes of different species varies and is only known for a few species (Inger et al., 2014). The duty cycle for PWM thus should be chosen that is unnoticeable for the suspects’ eyes. PWM can be constructed with inexpensive microcontrollers and resistors. We used an Arduino microcontroller following the instructions of Muhammad Aqib (https://create.arduino.cc/projecthub/muhammad-aqib/arduino-pwm-tutorial-ae9d71) to build a simple PMW controller. Depending on the operating voltage of the LEDs to use specific resistors need to be used.

Alternatively, we opted for a combination of filter-based dimming and analogous dimming by adjusting the voltage applied to the LEDs. This way, the drawback of analogous LED dimming (changes in the spectral characteristics) can be kept small. Neutral density filters reduce the incoming light regardless of spectral characteristics evenly by a certain percentage. We constructed a 3D printable dual filter revolver that can hold eight different neutral density filters (Fig 3, supporting information “Detailed Building Instructions”). Combining neutral density filters and altering voltages applied to the LED allowed us to adjust the emitted photons between 100 to 0.00001% continuously regardless of wavelength. When controlling LEDs via the applied voltage, it is of importance to monitor the emitted spectrum of each LED under these conditions since different voltages can alter their emission wavelength.

**Figure 3.**
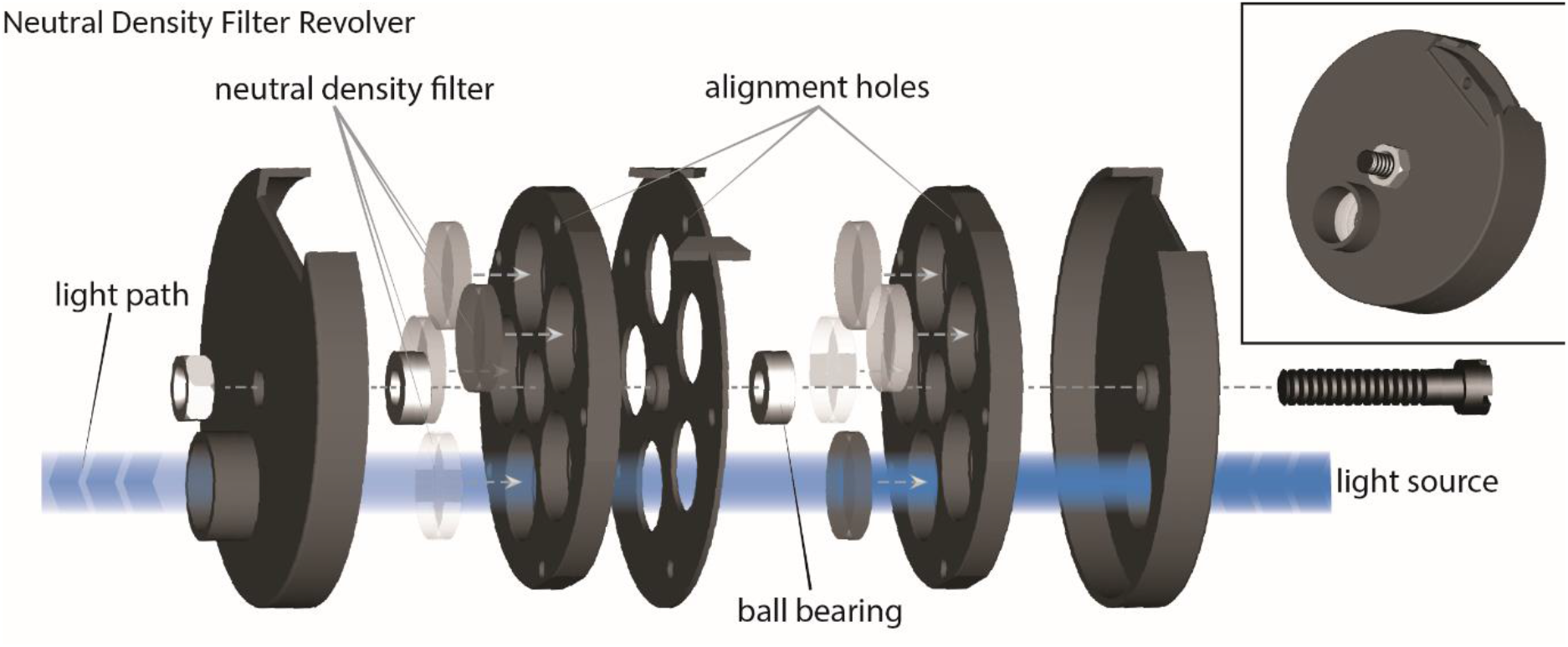
Modulating light. Neutral density filter revolver (NDR) allows to quickly changing different neutral density filters into the light path to influence the emitted light by the LED. The filters sit inside of two revolving wheels, each capable of holding up to 5 filters each. Alignments guarantee that the light path will not be partially blocked by intermediate positions of the filter.

Combining PWM and the described neutral density filter revolvers, finally, allows for the most accurate regulation of photon emission across broad ranges of light intensities.

#### Characterization of the LEDs

We recommend assessing the emitted spectrum of each LED before usage since the respective values given by the producers are not always accurate. For this purpose, we designed an adapter with which a LED light bulb can be put in a defined distance to a spectrometer. This way, it is also possible to measure the irradiance, which is a wavelength-independent quantity of light intensity. In a second step, both the spectrum and the intensity can be assessed for different applied voltages. We provide a simple script to convert irradiance into photon count (number of emitted photons), which is the most significant quantity for light intensity in photobiological experiments. By combining filter-based and analogous, voltage based dimming, the number of emitted photons can be adjusted to similar levels for different LEDs (Fig 4 A).

**Figure 4.**
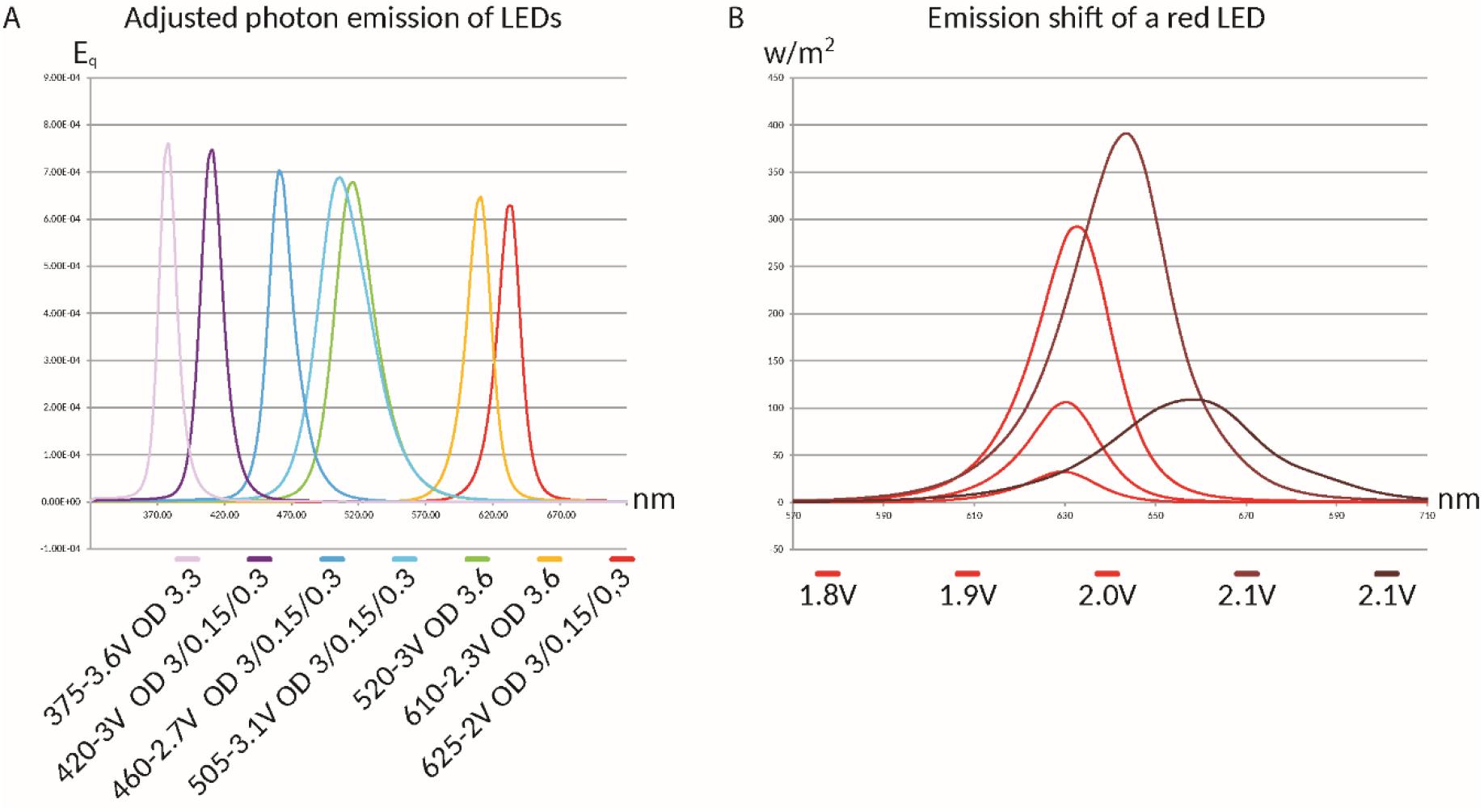
**A**. Using LEDs in combination with varying voltage and neutral density filters (optical density: OD) allows the regulation of photon emission to comparable levels. **B**. Special care should be given to regulating LEDs with voltages above their specification, as this can shift the emitted wavelength of the LED.

#### Light monitoring

Monitoring the exciting beam of light is of significant importance. In reality, not all LEDs are emitting exactly their advertised spectrum, and altering current can further shift their emitted spectrum (Fig 4 B). However, regardless of the light source used, it is also necessary to be able to make LEDs of different wavelengths comparable. Most commercially available products measure light intensity in LUX or Lumen that are both weighted based on human perception. To avoid human bias, we initially measured all our LEDs irradiance and then used a simple script to calculate the emitted photons of each LED. In conjunction with altering current and utilizing neutral density filter combinations, we were able to reduce the emitted photons of each LED to comparable levels (Fig 4 A).

#### Light Mixing

Light conditions in different depth differ significantly depending on time, weather conditions, seasons, and phytoplankton activity. UV radiation is usually absorbed within the first few meters, while blue light reaches the furthest within the water column. Furthermore, colored dissolved organic matter has a strong influence on the spectral characteristics of the water column by absorbing UV and blue light (DeGrandpre et al., 1996). Similarly, dissolved organic matter absorbs mainly short wavelengths. The spectral composition of light in the water column conveys important spatial and temporal information as well as other environmental conditions. Many planktonic animals come equipped with extraocular photoreceptors or additional eyes with yet unknown function and spectral sensitivity (DREWES & FOURTNER, 2007; Ohtsu, 1983; Ramirez et al., 2011) or may be able to perceive different wavelengths in the same eye (color vision). To be able to simulate very different light conditions, we conceived a modular light path of 3D printed parts, LEDs as the light source, and different dichroic mirrors to be able to combine different LEDs (Fig 2). Coupled with a beam splitter and a spectrometer, it allows us to adjust and monitor the emitted spectrum finely and thus control the stimulating light on the animals.

### Analysis Pipeline

#### TrackmateTaxis - tracking

The initial analysis pipeline starts with a Fiji (Schindelin et al., 2012) script that combines preprocessing of the videos as well as the tracking in a convenient way, Trackmatetaxis-tracking (Flowchart Fig 5, “Step-by-Step Guide” in supporting information). A GUI guides the user to all crucial steps and is capable of processing any number of videos autonomously. The image enhancements utilize build-in functions supplied by Fiji to calibrate the video, enhance contrast, and remove the background. The subsequent tracking utilizes Trackmate (Tinevez et al., 2017), a plugin originally designed for single-particle tracking. Trackers included in Trackmate fare well when it comes to identifying individual spots and even do well when spots are close to each other if settings have been adjusted to the animals. However, a significant problem of trackers is when two spots cross paths. Often the identity of spots can get lost during such events. Computer vision requires sophisticated algorithms to handle spots crossing over each other. To obtain statistically meaningful data sets in single recordings, we studied high numbers of animals and chose not to track the individuals over the entire duration. Instead, we recorded the cartesian coordinates of each detected spot, and only if the same spot was found in the next frame, we also let Trackmate calculate its velocity. If Trackmate is used via its own GUI, it will calculate all features of each spot and track. However, if it is executed using the provided script, the features that will be assessed are only those critical for subsequent analysis, and these are saved on disk as comma-separated values. This can save time when processing many recordings. The following analysis can work with files created either by the GUI of Trackmate or by the automatic script.

**Figure 5.**
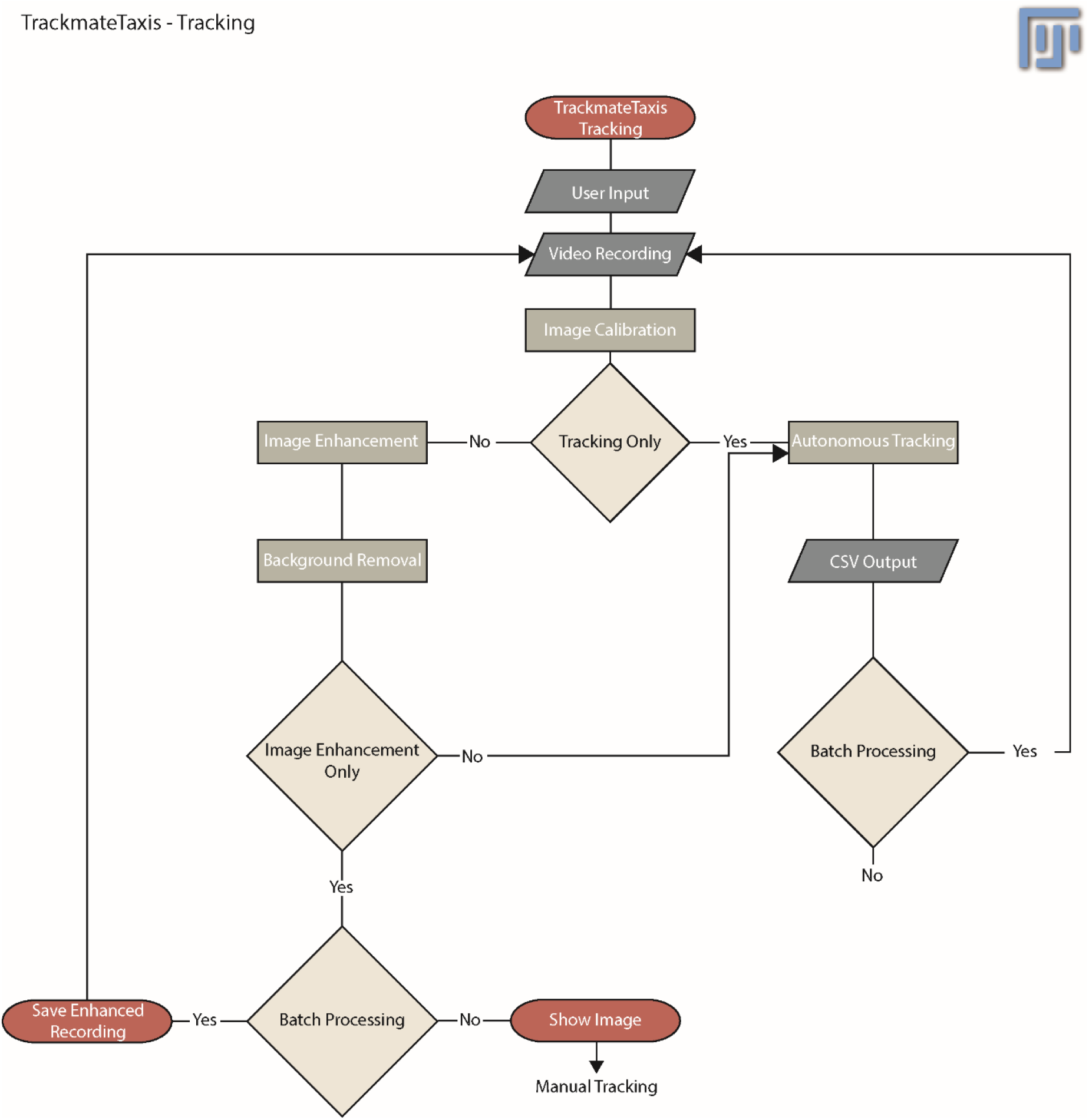
Flowchart of the software pipeline within Fiji. Diamonds are the decision by the user. Ovals denote beginning and ends. Rectangles are processing steps in the pipeline. A parallelogram depicts input and output. Initiating the trackmatetaxis script within Fiji will ask the user to specify working-directory and other parameters. The script fetches the avi-video files within the specified working directory and performs an image calibration. If the user decided to perform tracking only, all image enhancement steps are skipped, and the tracking with the plugin trackmate commences. If image enhancement has been chosen, the script will remove the background and increase the contrast to aid with the tracking. Usually, the script would pass the processed video to the tracker. If tracking is not desired, it will either save each file as a new video file or, if no batch processing has been chosen, will display the result directly within imageJ. This last step is supposed to aid in finding the best possible settings within Trackmate before the batch process can automatically work on all videos.

#### TrackmateTaxis – analyzing tracking data

The analysis of the tracked data can be done by the provided custom python script Trackmatetaxis-analysis (“Step-by-Step Guide” in Supporting information). It makes use of the CSV files created by Trackmate, has a GUI to ease usage, and is capable of processing many files autonomously. When starting a new experiment, the initial distribution of the animals in the cuvette may depend on many factors like gravity, chemical cues, pressure, swimming speed, or biorhythms. To make different experiments comparable, the script first calculates a baseline by averaging the tracked positions of all animals from frame “a” to “b” during the initial phase of the experiment. This baseline is subtracted from the median position 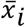 of a given frame i to achieve the normalized median position:

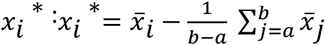

Furthermore, the script will also utilize the velocity information from the previous part of the pipeline to calculate the average speed of each frame. Velocity information can be particularly useful if the behavior of the animals is further impeded by chemical agents or other stimuli and can be used as a health indicator.

Similarly, the mean positions are calculated, and the script also adds 2%,9%,25%,50%,75%,91%, and 98% quantiles as well as minimum and maximum positions for each frame. To account for natural fluctuations of the animal’s responses, also time-averaged values will be saved in a new file, and a user-specified time interval of interest will create respective boxplots.

### Example Data

Two different animals were used with two different constellations of the setup. The annelid *M. fuliginosus* develops from pelagic planktotrophic larvae that remain within the water column for up to 36 days (Blake & Arnofsky, 1999). For this experiment, animals of an early-stage were used, which were fertilized and kept until day seven at 18C in a day/night cycle of 18/6 hours. This experiment was used to test the effect of different wavelengths on the animals. For this purpose, different LEDs were installed above a cuvette containing around 300 *M. fuliginosus* larvae. Every LED used was down-regulated to similar photon count emission with neutral density filters. Each recording started with 30 seconds of darkness to calculate the baseline and was followed by 15 seconds of illumination with different LEDs. The final 45 seconds were again without any light stimulus. Each video afterward was manually cropped just to show the cuvette containing the animals. Afterward, the Fiji script for tracking was initiated. A single video was used to perform image enhancement and find appropriate values for the detection. These values were then used to enhance and track the animals in all the recordings. Finally, the tracking analysis script was used to extract all positional data on the y-axis from 40 seconds to 42 seconds (corresponding to frames 3014 to 3161 @ 73.5fps). The resulting boxplot shows the median distribution for the animals for each wavelength during the previously specified time interval (Fig 6). The 7-day-old larvae of

**Figure 6.**
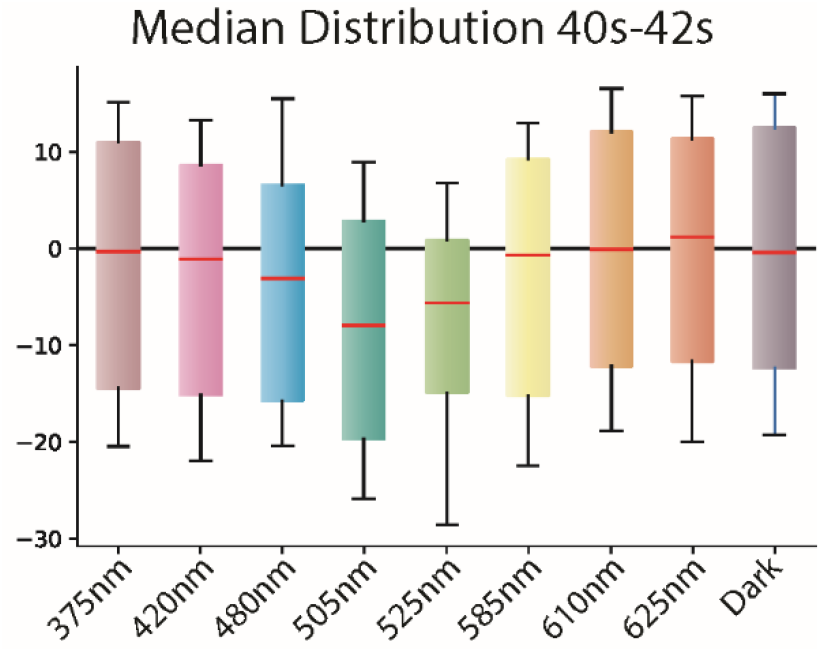
A boxplot showing the median distribution of the larvae of *Malacoceros fuliginosus* during illumination of different wavelengths. The animals show the greatest displacement towards the cyan wavelength.

Malacoceros prefer light in the turquoise part of the visible spectrum, showing a clear peak at 505nm. The displacement tapers off towards the yellow and ultraviolet parts of the spectrum until no reaction is detectable.

The second animal was the cosmopolitan bryozoan *Tricellaria inopinata*. The larvae are spherical and roughly measure 200μm in diameter. In contrast to *Malacoceros*, these larvae are lecithotrophic and only remain in the water column for approximately 24 hours. We transferred the larvae into a cuvette that had a UV LED of 355nm mounted underneath. The recording was a total of 54 seconds long with 27 seconds of darkness (corresponds to 3000 frames at 54.21817 frames per second), in the beginning, followed by 15 seconds illumination with the UV LED.

The remaining time was again in darkness. The videos were cropped to show only the cuvette, and then the Fiji script was used to enhance the video and track all the animals. The tracking result was analyzed, and the mean and median distribution of animals during each frame was calculated. The resulting curve demonstrates the median (Fig 7 A) and mean (Fig 7 B) distribution of the animals over the entire video. The median distribution tracked a swift change towards the source of illumination, the 75^th^ percentile reaches the edges of the vessel, and hence the displacement plateaus (Fig 7 A, arrow).

**Figure 7.**
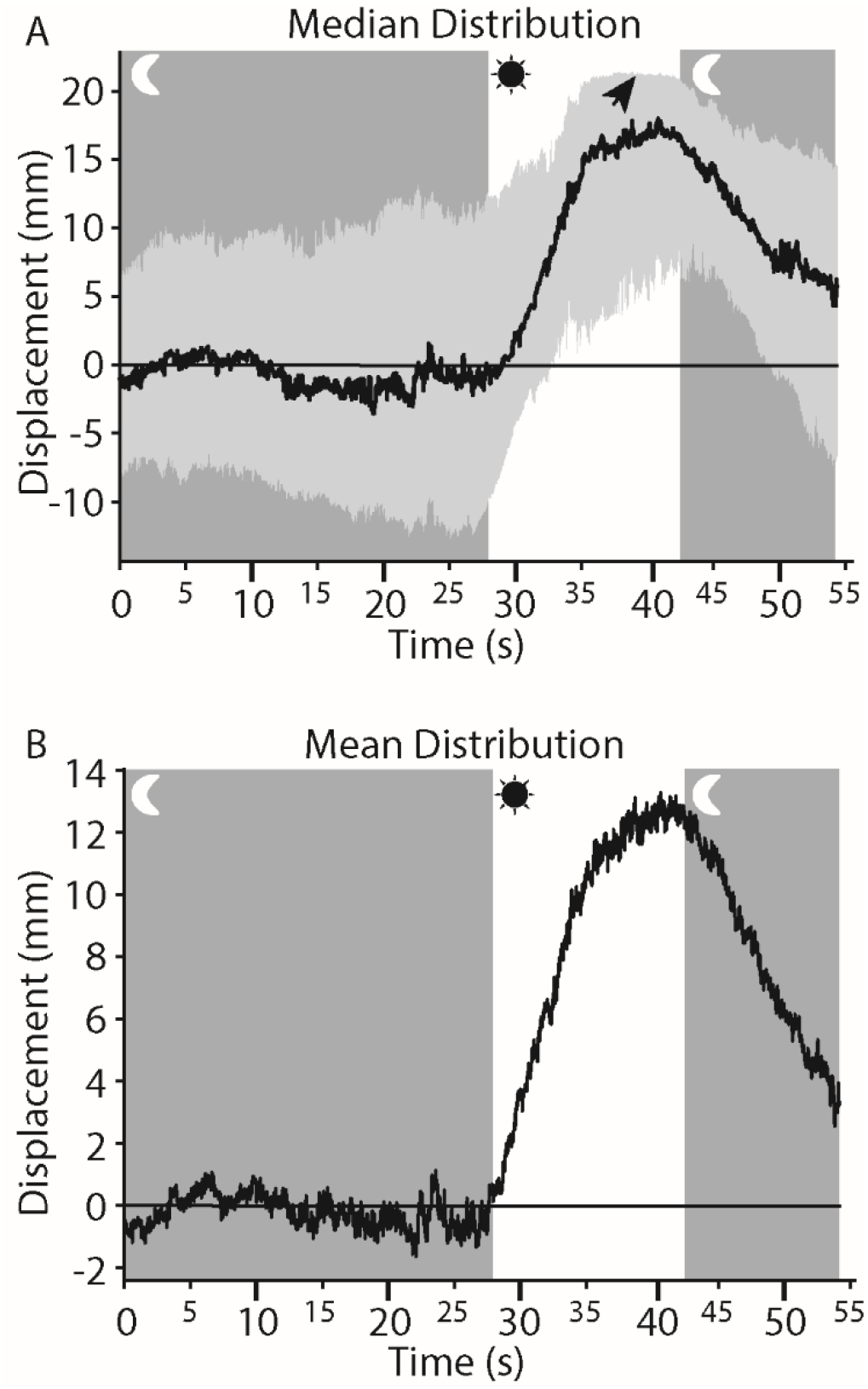
**A**. Median distribution of the larvae of *Tricellaria inopinata* showing their displacement towards a UV led over the entire duration of the recording. The light was turned on (white rectangle), and the animals started to swim towards the light source. Animals in the upper 75percentile hit the edge of the vessel, and the curve plateaus (arrow). **B**. The mean position of the same video as in **A**.

## Discussion

Quantification and reproducibility are prerequisites for proper analysis of behavioral data, be it in the context of behavioral biology, sensory biology, or ecology. In the case of macroscopic animals, observation and data analysis usually have to be adopted to a large extent to the specific research questions and the complexity of behavioral patterns observed. In the case of microscopic planktonic organisms, many questions can be addressed based on robust and comparable data on directed movements in the water column. We present an affordable and flexible experimental setup and an analysis pipeline, which can be used for this purpose and is applicable to study a wide range of organisms, irrespective of their specific swimming patterns.

With the help of 3D printing, our setup can easily be modified, extended, and adapted to more specific needs. The options are almost endless. The mounting options on our setup should allow the installation of many more add-ons like mechanical- or chemical stimuli. Furthermore, if 3D tracking is desirable, it would be possible to split the light path of one camera with a few 3D printed holders for mirrors, and the cuvette rotated by 45°. We hope that this setup with our analysis pipeline TrackmateTaxis will enable others also to conduct behavioral research on planktonic animals. All parts of this setup are designed with alterations in mind. It is not necessary to record videos with our setup to use our scripts and vice versa. As long as an avi-video is readable by Fiji, it can be processed. If a more advanced analysis is required, it is possible to either add to the existing script or use different ones. The behavioral spectrum of planktonic animals is as diverse as the animals itself and requires modular and flexible solutions that are easy to use and robust. This setup provides a solid foundation for future behavioral studies on a wide variety of planktonic animals.

## Supporting information

TrackmateTaxis Build Instructions

TrackmateTaxis Analysis Pipeline

## Acknowledgments

We thank Suman Kumar (University of Bergen) for providing *M. fuliginosus* larvae for the experiments. Børge Hamre (University of Bergen) supplied the Ramses Spectrometer, which we much appreciate.

## Authors contribution

CCD and HH designed the setup and the experiments. CCD created the 3D models, assembled the parts, wrote the software pipeline, and conducted the experiments. Both analyzed the data and wrote the manuscript. All authors gave final approval for the publication.

## Data availability

Data sets, scripts, building instructions, and 3D models will be made available at the repositories listed in table 1. **Respective Dois will be provided upon submission of the accepted manuscript**.

**Table 1.** Data availability for raw data, videos, and scripts.

| Data | Type of Data | Storage location | DOI |
| --- | --- | --- | --- |
| Behavioral Recordings of animals | Video (avi) | DataverseNO |  |
| Raw Analysis Data | Excel and CSV files | DataverseNO |  |
| ImageJ Plugin | Jython code | Github and Zenodo Archive |  |
| Python Analysis | Python code | Github and Zenodo Archive |  |
| Python SpectraRad to photon count | Python code | Github and Zenodo Archive |  |
| 3D Models | FCStd & Obj | DataverseNO |  |

## References

Blake, J. A., & Arnofsky, P. L. (1999). Reproduction and larval development of the spioniform polychaete with application to systematics and phylogeny. Hydrobiologia, 402, 57–106. https://doi.org/10.1023/A:1003784324125

Branson, K., Robie, A. A., Bender, J., Perona, P., & Dickinson, M. H. (2009). High-throughput ethomics in large groups of Drosophila. Nature Methods, 6(6), 451–457. https://doi.org/10.1038/nmeth.1328

Crispim Junior, C. F., Pederiva, C. N., Bose, R. C., Garcia, V. A., Lino-de-Oliveira, C., & Marino-Neto, J. (2012). ETHOWATCHER: Validation of a tool for behavioral and video-tracking analysis in laboratory animals. Computers in Biology and Medicine, 42(2), 257–264. https://doi.org/10.1016/j.compbiomed.2011.12.002

DeGrandpre, M. D., Vodacek, A., Nelson, R. K., Bruce, E. J., & Blough, N. V. (1996). Seasonal seawater optical properties of the U.S. Middle Atlantic Bight. Journal of Geophysical Research: Oceans, 101(C10), 22727–22736. https://doi.org/10.1029/96JC01572

Drewes, C. D., & Fourtner, C. R. (2007). Hindsight and Rapid Escape in a Freshwater Oligochaete. The Biological Bulletin, 177(3), 363–371. https://doi.org/10.2307/1541596

Effertz, C., & von Elert, E. (2014). Light intensity controls anti-predator defences in Daphnia: the suppression of life-history changes. Proceedings of the Royal Society B: Biological Sciences, 281(1782), 20133250–20133250. https://doi.org/10.1098/rspb.2013.3250

Elliott, G. R. D., MacDonald, T. A., & Leys, S. P. (2004). Sponge Larval Phototaxis: A Comparative Study. In Bollettino dei Musei e degli Istituti Biologici (Vol. 68).

Erkal, J. L., Selimovic, A., Gross, B. C., Lockwood, S. Y., Walton, E. L., McNamara, S., Martin, R. S., & Spence, D. M. (2014). 3D printed microfluidic devices with integrated versatile and reusable electrodes. Lab Chip, 14(12), 2023–2032. https://doi.org/10.1039/C4LC00171K

Fernö, A., Huse, I., Juell, J. E., & Bjordal, Å. (1995). Vertical distribution of Atlantic salmon (Salmo solar L.) in net pens: trade-off between surface light avoidance and food attraction. Aquaculture, 132(3–4), 285–296. https://doi.org/10.1016/0044-8486(94)00384-Z

Fields, D. M., & Yen, J. (1997). The escape behavior of marine copepods in response to a quantifiable fluid mechanical disturbance. Journal of Plankton Research, 19(9), 1289–1304. https://doi.org/10.1093/plankt/19.9.1289

Gal, G., Loew, E. R., Rudstam, L. G., & Mohammadian, a M. (1999). Light and diel vertical migration: spectral sensitivity and light avoidance by Mysis relicta. Canadian Journal of Fisheries and Aquatic Sciences, 56(1975), 311–322. https://doi.org/2

Gallager, S. M., Yamazaki, H., & Davis, C. S. (2004). Contribution of fine-scale vertical structure and swimming behavior to formation of plankton layers on Georges Bank. Marine Ecology Progress Series, 267, 27–43. https://doi.org/10.3354/meps267027

Garm, A., O’Connor, M., Parkefelt, L., & Nilsson, D.-E. (2007). Visually guided obstacle avoidance in the box jellyfish Tripedalia cystophora and Chiropsella bronzie. Journal of Experimental Biology, 210(20), 3616–3623. https://doi.org/10.1242/jeb.004044

Hart, A. (2006). Behavior. WormBook, 1–67. https://doi.org/10.1895/wormbook.1.87.1

Hays, G. C. (2003). A review of the adaptive significance and ecosystem consequences of zooplankton diel vertical migrations. Hydrobiologia, 503, 163–170. https://doi.org/10.1023/B:HYDR.0000008476.23617.b0

Inger, R., Bennie, J., Davies, T. W., & Gaston, K. J. (2014). Potential biological and ecological effects of flickering artificial light. PLoS ONE, 9(5). https://doi.org/10.1371/journal.pone.0098631

Jékely, G., Colombelli, J., Hausen, H., Guy, K., Stelzer, E., Nédélec, F., & Arendt, D. (2008). Mechanism of phototaxis in marine zooplankton. Nature, 456(7220), 395–399. https://doi.org/10.1038/nature07590

Kiørboe, T. (2007). Mate finding, mating, and population dynamics in a planktonic copepod Oithona davisae: There are too few males. Limnology and Oceanography, 52(4), 1511–1522. https://doi.org/10.4319/lo.2007.52.4.1511

Marsden, J. R. (1986). Response to light by trochophore larvae of Spirobranchus giganteus. Marine Biology, 93(1), 13–16. https://doi.org/10.1007/BF00428649

Miller, S. E., & Hadfield, M. G. (1986). Ontogeny of phototaxis and metamorphic competence in larvae of the nudibranch Phestilla sibogae Bergh (Gastropoda : Opisthobranchia). Journal of Experimental Marine Biology and Ecology, 97(1), 95–112. https://doi.org/10.1016/0022-0981(86)90070-5

Ohtsu, K. (1983). UV–visible antagonism in extraocular photosensitive neurons of the anthomedusa, spirocodon saltatrix (tilesius). Journal of Neurobiology, 14(2), 145–155. https://doi.org/10.1002/neu.480140206

Oskarsson, M., Kjellberg, T., Palmér, T., Nilsson, D.-E., & Åström, K. (2015). Visual Tracking of Box Jellyfish. In Computer Vision and Pattern Recognition in Environmental Informatics (pp. 107–122). https://doi.org/10.4018/978-1-4666-9435-4.ch006

Pérez-Escudero, A., Vicente-Page, J., Hinz, R. C., Arganda, S., & De Polavieja, G. G. (2014). IdTracker: Tracking individuals in a group by automatic identification of unmarked animals. Nature Methods, 11(7), 743–748. https://doi.org/10.1038/nmeth.2994

Petie, R., Garm, A., & Nilsson, D.-E. (2011). Visual control of steering in the box jellyfish Tripedalia cystophora. Journal of Experimental Biology, 214(17), 2809–2815. https://doi.org/10.1242/jeb.057190

Piraino, S., Zega, G., Di Benedetto, C., Leone, A., Dell’Anna, A., Pennati, R., Candia Carnevali, D., Schmid, V., & Reichert, H. (2011). Complex neural architecture in the diploblastic larva of Clava multicornis (Hydrozoa, Cnidaria). The Journal of Comparative Neurology, 519(10), 1931–1951. https://doi.org/10.1002/cne.22614

Ramirez, M. D., Speiser, D. I., Pankey, M. S., & Oakley, T. H. (2011). Understanding the dermal light sense in the context of integrative photoreceptor cell biology. Visual Neuroscience, 28(04), 265–279. https://doi.org/10.1017/s0952523811000150

Rawat, S., Komatsu, S., Markman, A., Anand, A., & Javidi, B. (2017). Compact and field-portable 3D printed shearing digital holographic microscope for automated cell identification. Applied Optics, 56(9), D127. https://doi.org/10.1364/AO.56.00D127

Ringelberg, J. (1964). The positively phototactic reaction of daphnia magna straus: A contribution to the understanding of diurnal vertical migration. Netherlands Journal of Sea Research, 2(3), 319–406. https://doi.org/10.1016/0077-7579(64)90001-8

Rodríguez, Á., Bermúdez, M., Rabuñal, J. R., & Puertas, J. (2015). Fish tracking in vertical slot fishways using computer vision techniques. Journal of Hydroinformatics, 17(2), 275. https://doi.org/10.2166/hydro.2014.034

Rodriguez, A., Zhang, H., Klaminder, J., Brodin, T., Andersson, P. L., & Andersson, M. (2017). ToxTrac: A fast and robust software for tracking organisms. Methods in Ecology and Evolution, 2018(July 2017), 460–464. https://doi.org/10.1111/2041-210X.12874

Saberioon, M. M., & Cisar, P. (2016). Automated multiple fish tracking in three-Dimension using a Structured Light Sensor. Computers and Electronics in Agriculture, 121, 215–221. https://doi.org/10.1016/j.compag.2015.12.014

Samson, A. L., Ju, L., Kim, H. A., Zhang, S. R., Lee, J. A. A., Sturgeon, S. A., Sobey, C. G., Jackson, S. P., & Schoenwaelder, S. M. (2015). MouseMove: An open source program for semi-automated analysis of movement and cognitive testing in rodents. Scientific Reports, 5(May), 1–11. https://doi.org/10.1038/srep16171

Schindelin, J., Arganda-Carreras, I., Frise, E., Kaynig, V., Longair, M., Pietzsch, T., Preibisch, S., Rueden, C., Saalfeld, S., Schmid, B., Tinevez, J. Y., White, D. J., Hartenstein, V., Eliceiri, K., Tomancak, P., & Cardona, A. (2012). Fiji: An open-source platform for biological-image analysis. Nature Methods, 9(7), 676–682. https://doi.org/10.1038/nmeth.2019

Swift, M. C., & Forward, R. B. (1980). Photoresponses of Chaoborus larvae. Journal of Insect Physiology, 26(6), 365–371. https://doi.org/10.1016/0022-1910(80)90006-2

Symes, M. D., Kitson, P. J., Yan, J., Richmond, C. J., Cooper, G. J. T., Bowman, R. W., Vilbrandt, T., & Cronin, L. (2012). Integrated 3D-printed reactionware for chemical synthesis and analysis. Nature Chemistry, 4(5), 349–354. https://doi.org/10.1038/nchem.1313

Thorson, G. (1964). Light as an ecological factor in the dispersal and settlement of larvae of marine bottom invertebrates. Ophelia, 1(1), 167–208. https://doi.org/10.1080/00785326.1964.10416277

Tinevez, J. Y., Perry, N., Schindelin, J., Hoopes, G. M., Reynolds, G. D., Laplantine, E., Bednarek, S. Y., Shorte, S. L., & Eliceiri, K. W. (2017). TrackMate: An open and extensible platform for single-particle tracking. Methods, 115, 80–90. https://doi.org/10.1016/j.ymeth.2016.09.016

Tranter, D. J., Bulleid, N. C., Campbell, R., Higgins, H. W., Rowe, F., Tranter, H. A., & Smith, D. F. (1981). Nocturnal movements of phototactic zooplankton in shallow waters. Marine Biology, 61(4), 317–326. https://doi.org/10.1007/BF00401571

Veraszto, C., Guhmann, M., Jia, H., Rajan, V. B. V., Bezares-Calderon, L. A., Lopez, C. P., Randel, N., Shahidi, R. R., Michiels, N. K., Yokoyama, S., Tessmar-Raible, K., Jekely, G., Verasztó, C., Gühmann, M., Jia, H., Babu Veedin Rajan, V., Bezares-Calderón, L. A., Piñeiro Lopez, C., Randel, N., … Jékely, G. (2018). Ciliary and rhabdomeric photoreceptor-cell circuits form a spectral depth gauge in marine zooplankton. BioRxiv, 1–26. https://doi.org/10.1093/femsyr/foy001/4794945

Xiang, Y., Yuan, Q., Vogt, N., Looger, L. L., Jan, L. Y., & Jan, Y. N. (2010). Light-avoidance-mediating photoreceptors tile the Drosophila larval body wall. Nature, 468(7326), 921–926. https://doi.org/10.1038/nature09576

