## Supplementary material for "3D-printed Planktonic Observational Setup and Analysis Pipeline TrackmateTaxis": TrackmateTaxis Build Instructions

### TrackmateTaxis Setup
### Building Instructions

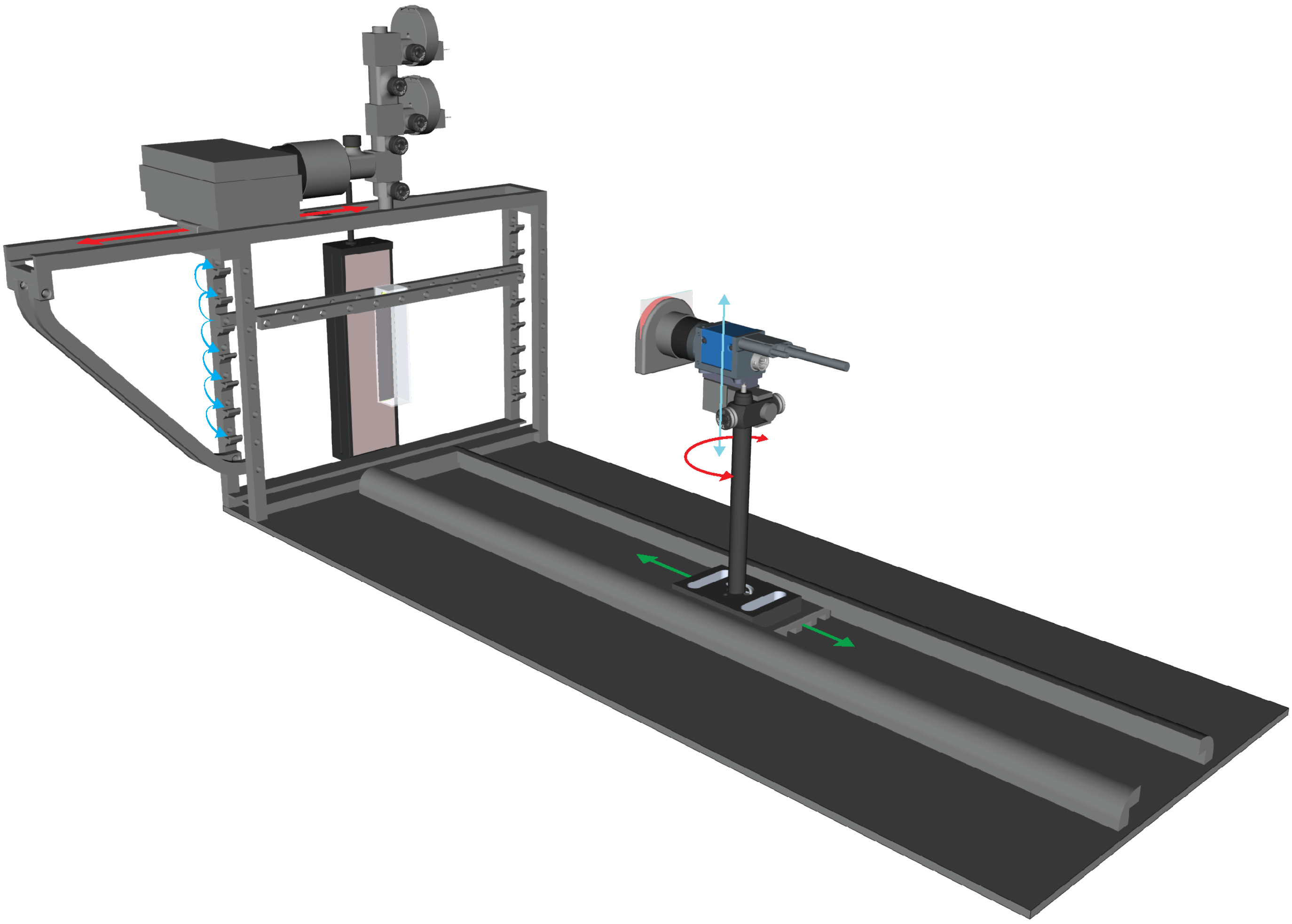

### General notes on construction and assembly

- Most of the illustrations are in a form of exploded diagrams to better illustrate the parts that they contain and how they can be assembled. An assembled view is usually in one of the corners.

- 3D printed part models were created with FreeCad, which is a free CAD software. The files can be edited in any way desired.

- 3D printed parts do depend on the printer/printing method used. Some prints might warp, shrink or otherwise differ from the computer model! Every printer is different and tolerances and minimum requirements differ.

- Some parts require filing to fit perfectly into each other.

- Screw slots have no thread printed. Most times the thread will be created by screwing unless for the Connector pieces (see Additional Printed Parts pages 6/7 and Light Mixer pages 8/9).

**Tools needed:**

- File/Sandpaper - to ensure a tight fit some parts need filing
- Glue - Some parts need to be glued together
- Scews - M4,M5 and M6 scews are needed including bolts and washers
- Baseplate - A plate to mount the setup on (300mm x 760mm)

**Light path construction:**

- This paper gives 3 options for a light path. From very simple to rather complex.
- The parts are all interchangable between these setups.
- There is an alternativ neutral density holder instead of the revolver that is simpler.
- To alter the length of the light path use different length of ø12mm or ø16mm tubing.
- The Connector pieces stabilize the light path better than just ø16mm tubing.

**LEDs:**

All LEDs bought from pur-led.de

| Wavelength (nm) | Part Number |
|---|---|
| 375 | 414375030 |
| 420 | 4140501 |
| 460 | 92501 |
| 500 | 2650501 |
| 520 | 50501 |
| 585 | 630501 |
| 610 | 6990501 |
| 625 | 80501 |

### General overview

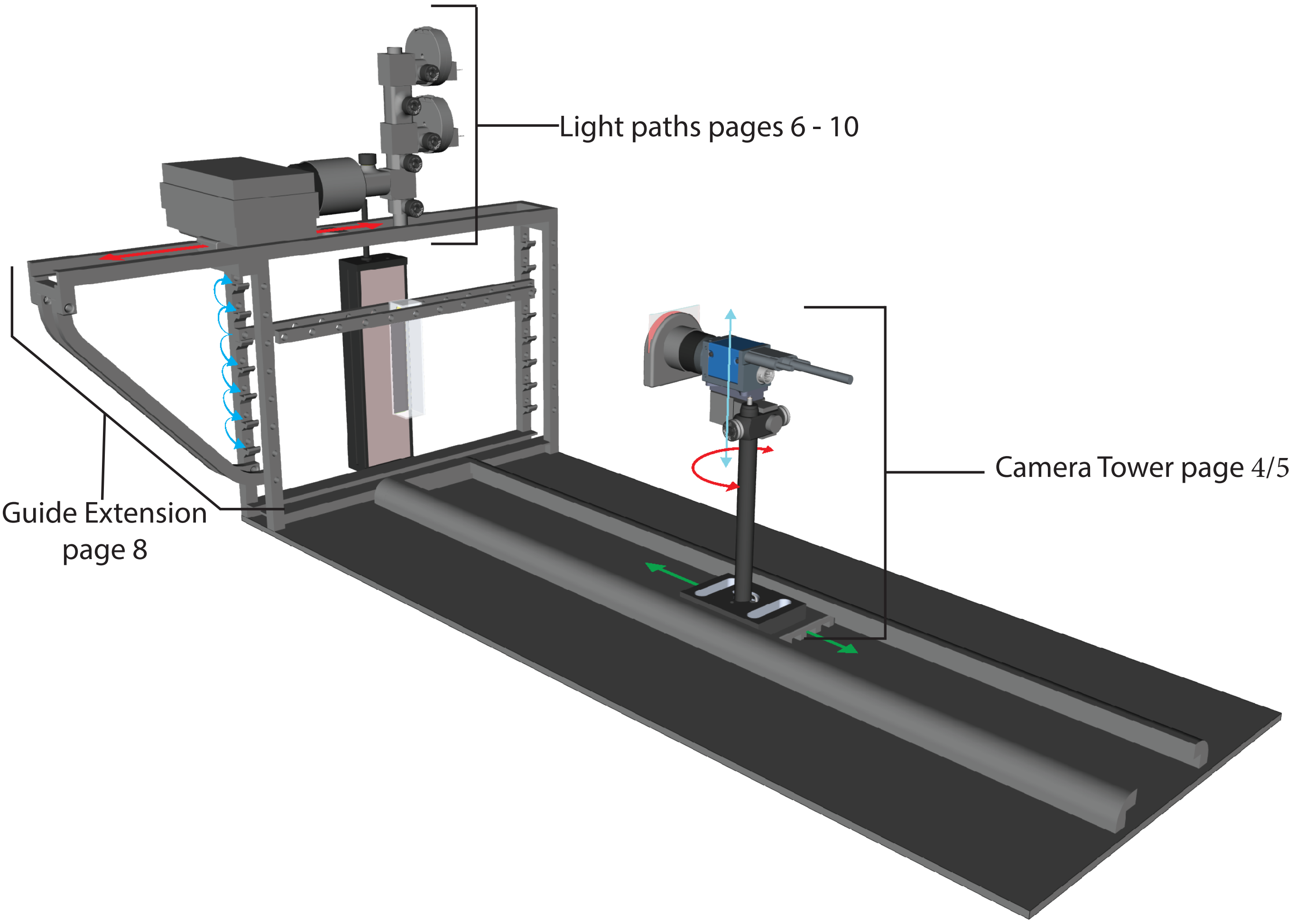

Contact Information
If you need any additional info, do not hesitate to contact me.

ccdoering

/Clemens_Doering

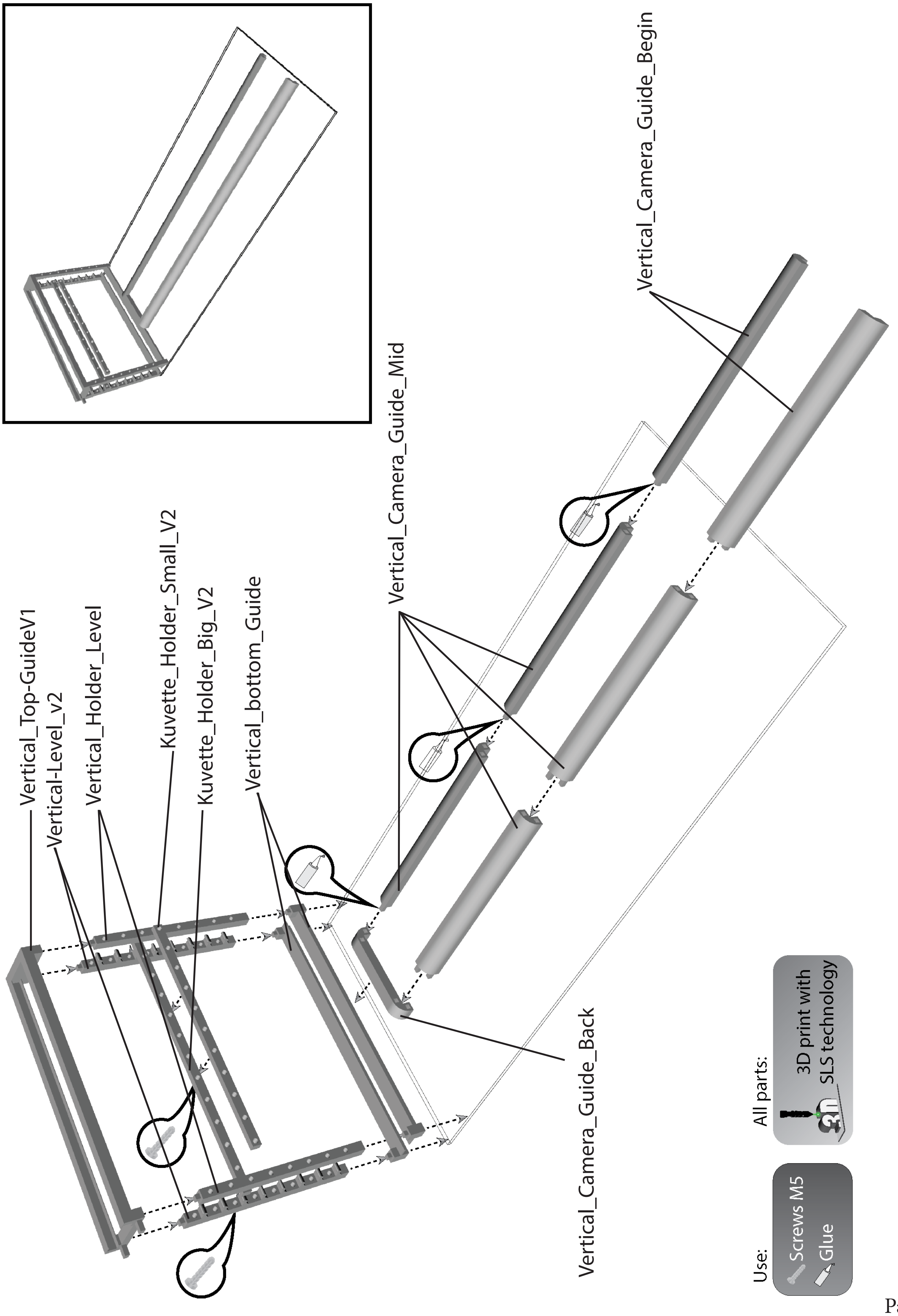
Vertical_Top-GuideV1
Vertical-Level_v2
Vertical_Holder_Level
Kuvette_Holder_Small_V2
Kuvette_Holder_Big_V2
Vertical_bottom_Guide
Vertical_Camera_Guide_Mid
Vertical_Camera_Guide_Begin
Vertical_Camera_Guide_Back
Vertical_Camera_Guide_Back
Use:
Screws M5
Glue
All parts:
3D print with
SLS technology

### Camera Tower

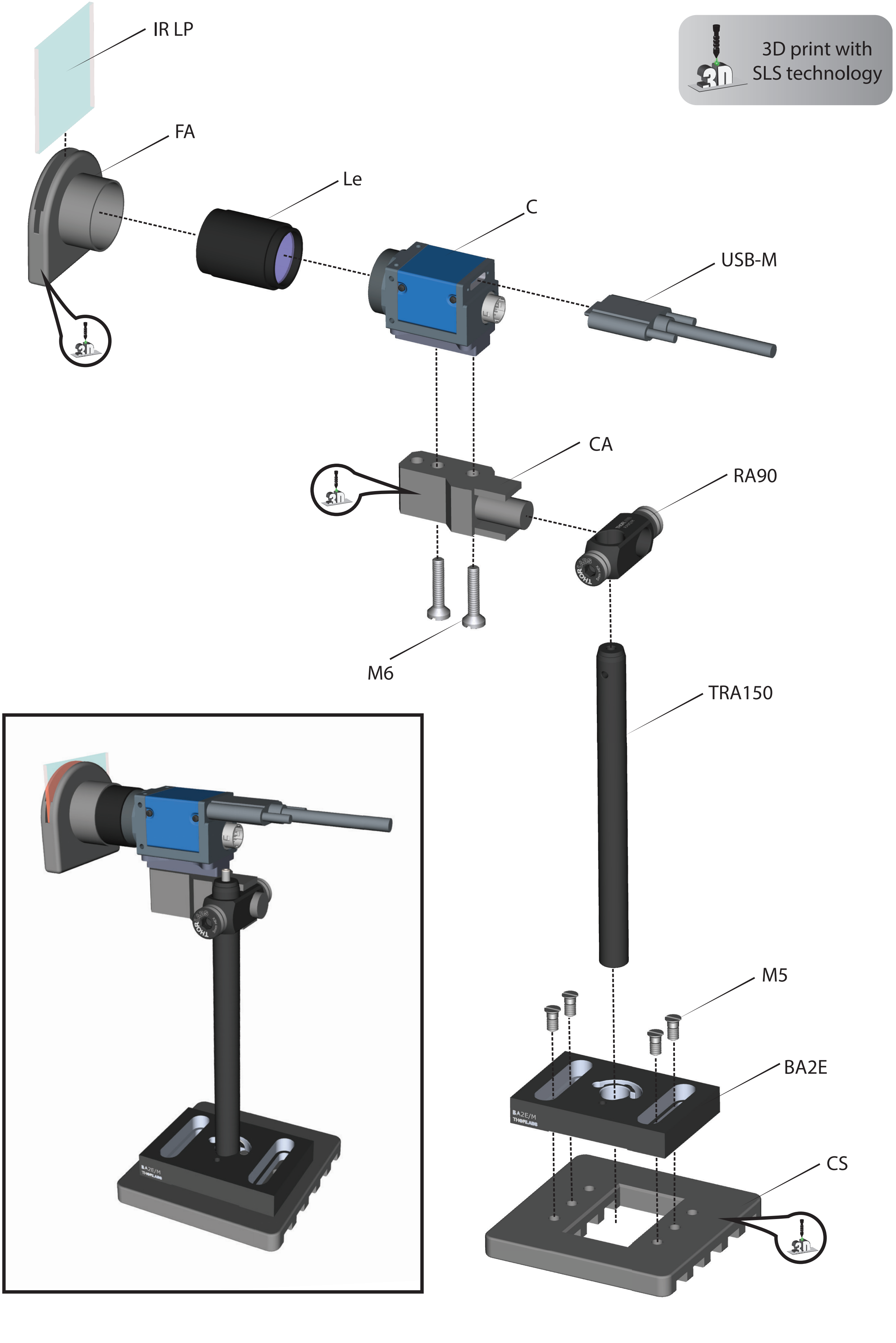

### Camera Tower Notes

##### 3D Printed Parts:

| Abbr | Name | 3D Model name |
|---|---|---|
| FA | Filter Adapter | Filter_Adapter |
| CA | Camera Adapter | Camera_Adapter |
| CS | Camera Slider | Camera_Slider |

##### Other Parts

| Abbr | Name | Details |
|---|---|---|
| IR LP | IR Longpass Filter | Reichmann Feinoptik RG610 50mm x 50mmx 2mm |
| Le | Lens | Ricoh FL-CC1614-2M |
| C | Camera | Imaging Source DMK 23UX236 |
| RA90 | Right-Angle Clamp | Thorlabs RA90 |
| TRA150 | TRA150 | Thorlabs Optical Post TRA150 |
| BA2E | Base | Thorlabs base for optical posts BA2E |

##### Notes

The Camera Adapter also exists in an unleveld version to introduce another axis of rotation.
See page 6/7 - Additional Printed Parts -> Rotatable Camera Adapter

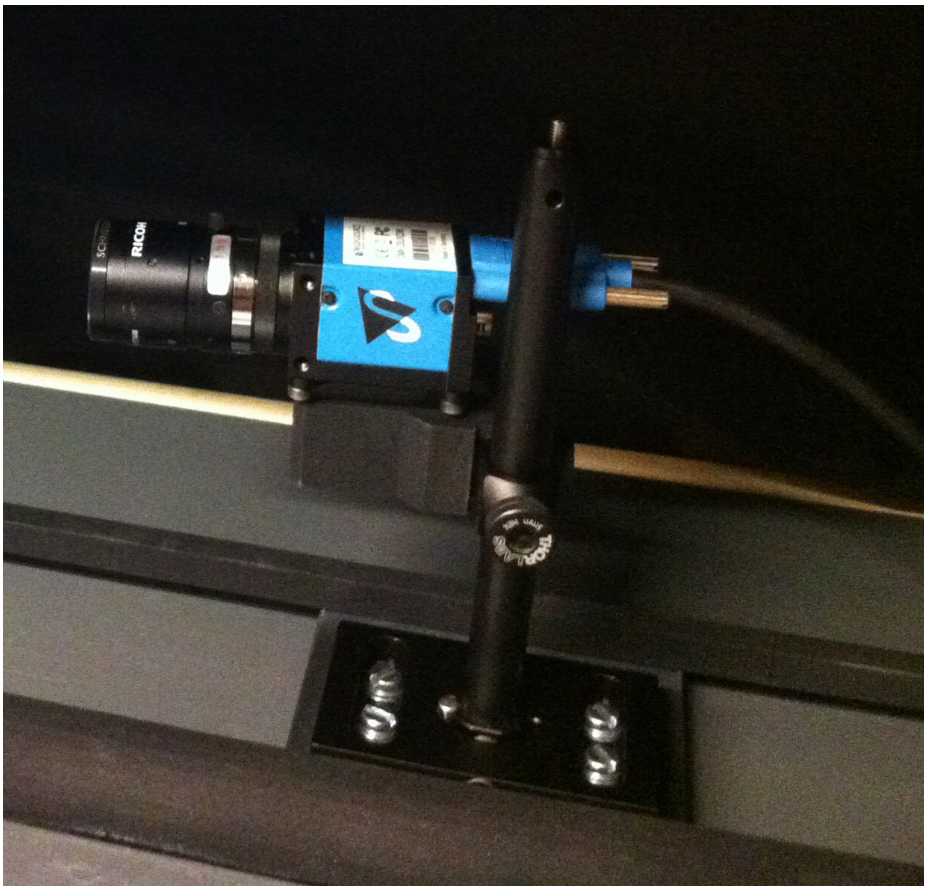

Example camera tower in the setup without longpass filter

#### Neutral Density Filter Revolver (NDR)

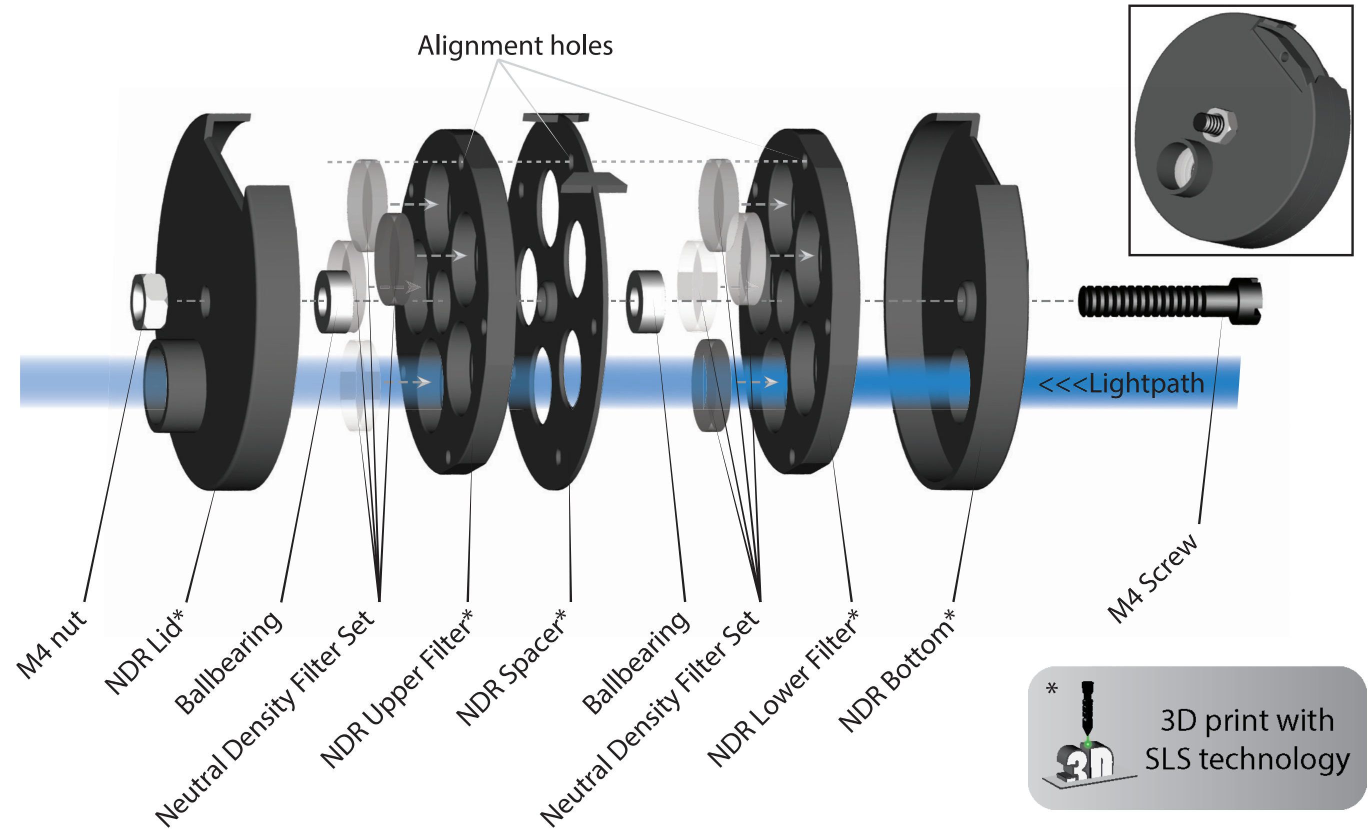

#### Bottom NDR Light Path

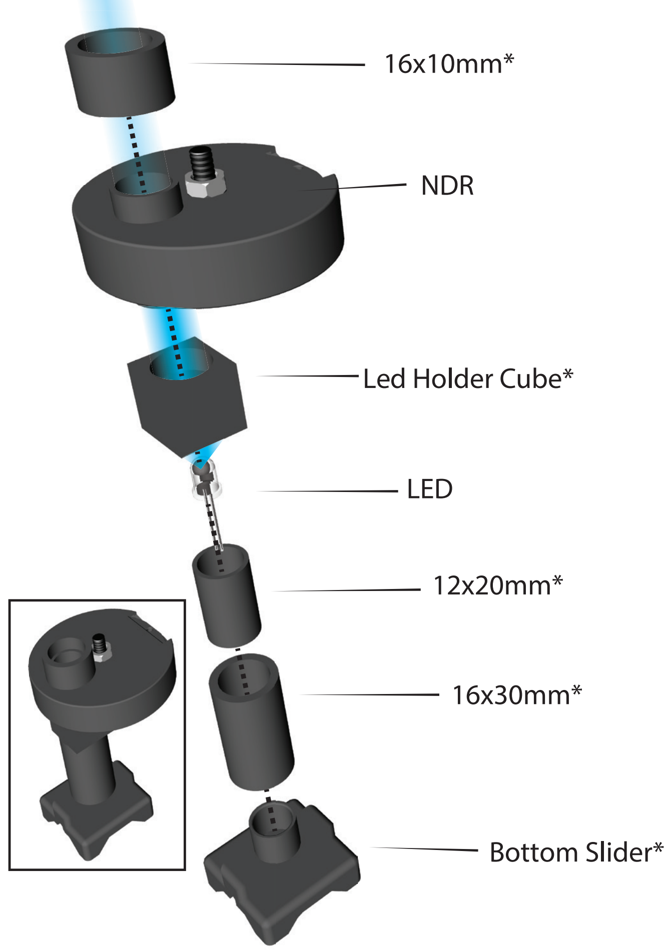

#### Additional Printed Parts

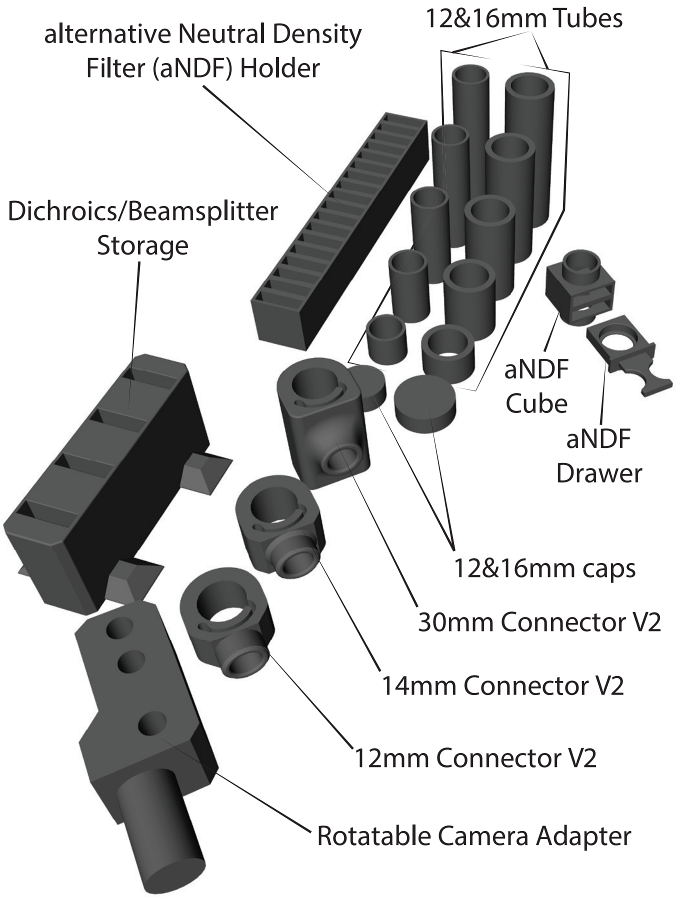

#### Neutral Density Filter Revolver Parts Notes

**3D Printed Parts:**

| Name | 3D Model name |
|---|---|
| NDR Lid | NDR_Lid |
| NDR Upper Filter | NDR_Upper_Filter |
| NDR Spacer | NDR_Spacer |
| NDR Lower Filter | NDR_Lower_Filter |
| NDR Bottom | NDR_Bottom |

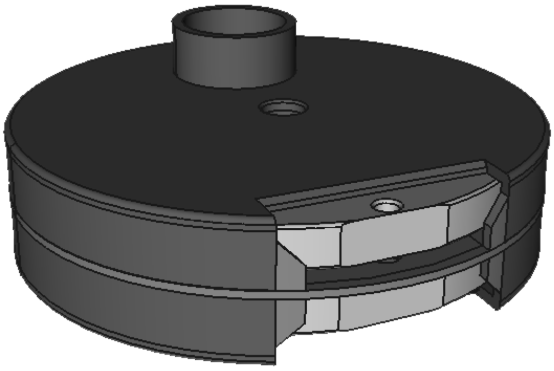

**Other Parts:**

| Name | Details |
|---|---|
| Ballbearing | ø = 8mm, inside ø = 4mm, RS Components 612-5767 |
| Neutral Density Filter Set | ø = 10mm, Reichmann Feinoptik (page 10 for details) |

**Notes**

- To align filters, use a dulled needle/nail and place into aligment holes
- Each side of the filter has a flattened part on the edge to allow place for a label
- Leave one filter slot empty in case no filter is needed

#### Bottom NDR Light Path Notes

**3D Printed Parts:**

| Name | 3D Model name |
|---|---|
| 16x10mm | 16mm-x-10mm_Tube |
| LED Holder Cube | LED_Holder_Cube |
| 12x20mm | 12mm-x-20mm_Tube |
| 16x30mm | 16mm-x-30mm_Tube |
| Bottom Slider | Bottom_Slider |

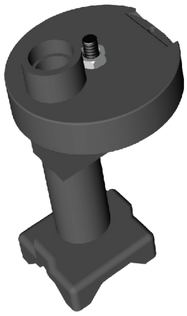

**Other Parts:**

| Name | Details |
|---|---|
| LED | See general notes on page 1 |
| NDR | See above on this page |

#### Additional Printed Parts Notes

**3D Printed Parts:**

| Name | 3D Model name |
|---|---|
| aNDF Holder | aNDF_Holder |
| aNDF Cube | aNDF_Cube |
| aNDF Drawer | aNDF_Drawer |
| 12/16mm Tubes | 12/16mm-x-[LENGTH]mm_Tube |
| 12/16mm Cap | 12/16mm_Cap |
| Connector V2 | 16mm_Connector_[LENGTH]mm_v3* |
| Rotatable Camera Adapter | Rotatable_Camera_Adapter |
| Dichroic/Beamsplitter Storage | Mirror_Storage_v2 |

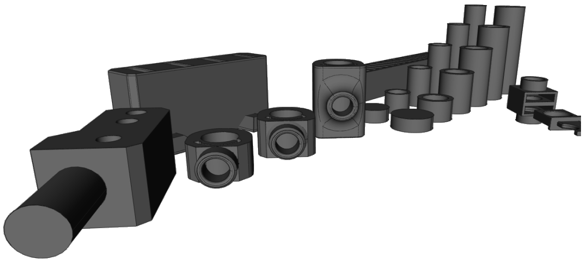

* Assembly of Connector see page 8/9 Light Mixer "Notes **"

### Light Mixer

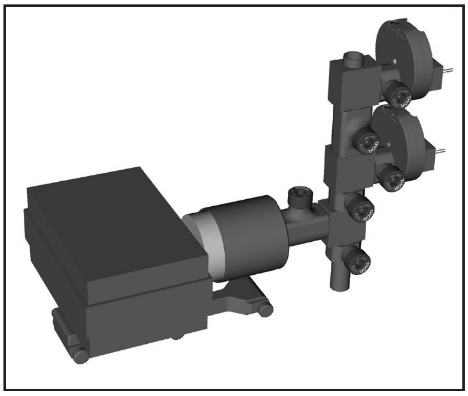

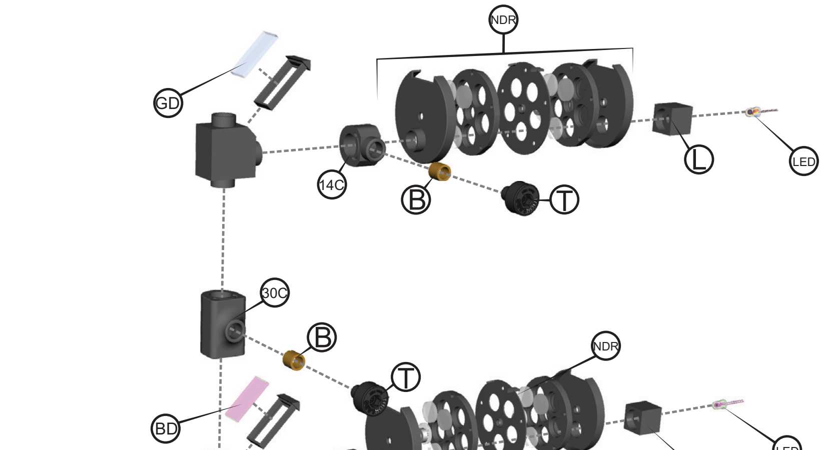

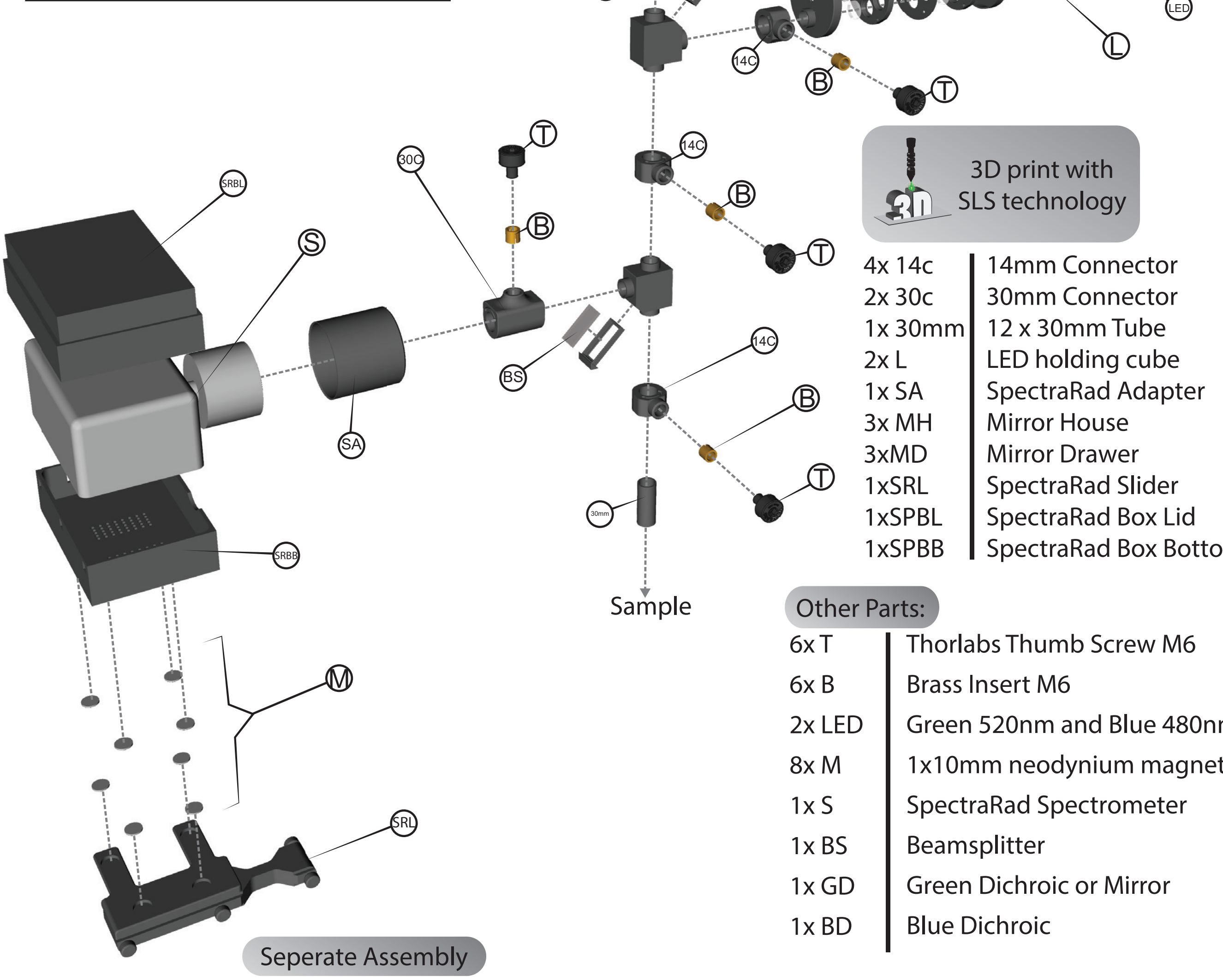

3D print with SLS technology

| | |
|---|---|
| 4x 14c | 14mm Connector |
| 2x 30c | 30mm Connector |
| 1x 30mm | 12 x 30mm Tube |
| 2x L | LED holding cube |
| 1x SA | SpectraRad Adapter |
| 3x MH | Mirror House |
| 3xMD | Mirror Drawer |
| 1xSRL | SpectraRad Slider |
| 1xSPBL | SpectraRad Box Lid |
| 1xSPBB | SpectraRad Box Bottom |

Other Parts:

| | |
|---|---|
| 6x T | Thorlabs Thumb Screw M6 |
| 6x B | Brass Insert M6 |
| 2x LED | Green 520nm and Blue 480nm |
| 8x M | 1x10mm neodynium magnet |
| 1x S | SpectraRad Spectrometer |
| 1x BS | Beamsplitter |
| 1x GD | Green Dichroic or Mirror |
| 1x BD | Blue Dichroic |

Seperate Assembly

2x NDR | Neutral Density Revolver consits of printed and other parts

### Guide Extension

This extension supports the light mixer and adds flexibility to its position.
It consits of 3 printed parts and needs 4 m5 screws and bolts.

3D Model names

Guide_Extension

Guide_Extension_Support_Beam

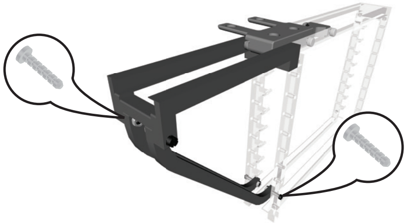

**3D Printed Parts:**

| Abbr | Name | 3D Model name |
|---|---|---|
| 14C | 14mm Connector | 16mm_Connector_14mm_v3 |
| 30C | 30mm Connector | 16mm_Connector_30mm_v3 |
| 30mm | 30mm Tube | 12mm-x-30mm_Tube |
| L | LED Holding Cube | LED_Holding_Cube |
| SA | SpectraRad Adapter | SpectraRad_Adapter |
| MH | Mirror House | Mirror_House* |
| MD | Mirror Drawer | Mirror_Drawer |
| SRL | SpectraRad Slider | SpectraRad-Slider |
| SRBT | SpectraRad Box Top | SpectraRad-Box-Lid |
| SRBB | SpectraRad Box Bottom | SpectraRad-Box-Bottom |

**Other Parts:**

| Abbr | Name | Details |
|---|---|---|
| T | Thumbscrew | Thorlabs Thumbscrew TS6H/M |
| B | Brass Inserts | M6 ø = 8mm, depth = 12.7mm (RS Components 278-578)** |
| LED | LEDs | See general notes page 1 "Light Paths" |
| M | Magnet | Neodynium magnet 1x10mm*** |
| S | Spectrometer | SpectraRad B&WTek inc. BSR112E-VIS/NIR |
| BS | Beamsplitter | Omega Optical 12x26mm 201279, see page 11 |
| GD | Green Dichroic | Omega Optical 12x26mm 565DRLPXR 201278, see page 11 |
| BD | Blue Dichroic | Omega Optical 12x26mm 505DRLP 201277, see page 11 |
| Mir | Mirror | Omega Optical 12x26mm 201280, see page 11 |

**Notes**

* Also available with either Blue, Green, UV, Mirror, or BS printed on the side (Mirror_House-Blue, -Green, -UV, -Mirror, and -BS as 3D model names)
** Assembly of Connector pieces! The brass inserts need to be shortened to about 8mm before they are installed into the opening of the connector piece.
*** Magnets from RS components: 811-2640 and/or 811-2631. The bottom slider can easily hold two magnets, increasing the needed magnets to 12 for extra supporting strength.

**Construction Tips**

When using Longpass filters, the Light Mixer should be constructed in such a way, that the LED with the longer wavelengths are the furthest away from the sample.
Example Yellow, Green, Blue, UV -> sample.
By adding standart mirrors it is possible to add any LED of longer wavelength than the existing dichroics. This means, in the example on the other page with two LEDs, it is not necessary to have two dichroics. The green dichroic can be replaced by a mirror, too.
The mirrors/dichroics describe here allow mixing of up to 4 different LEDs.

#### Basic Light Path

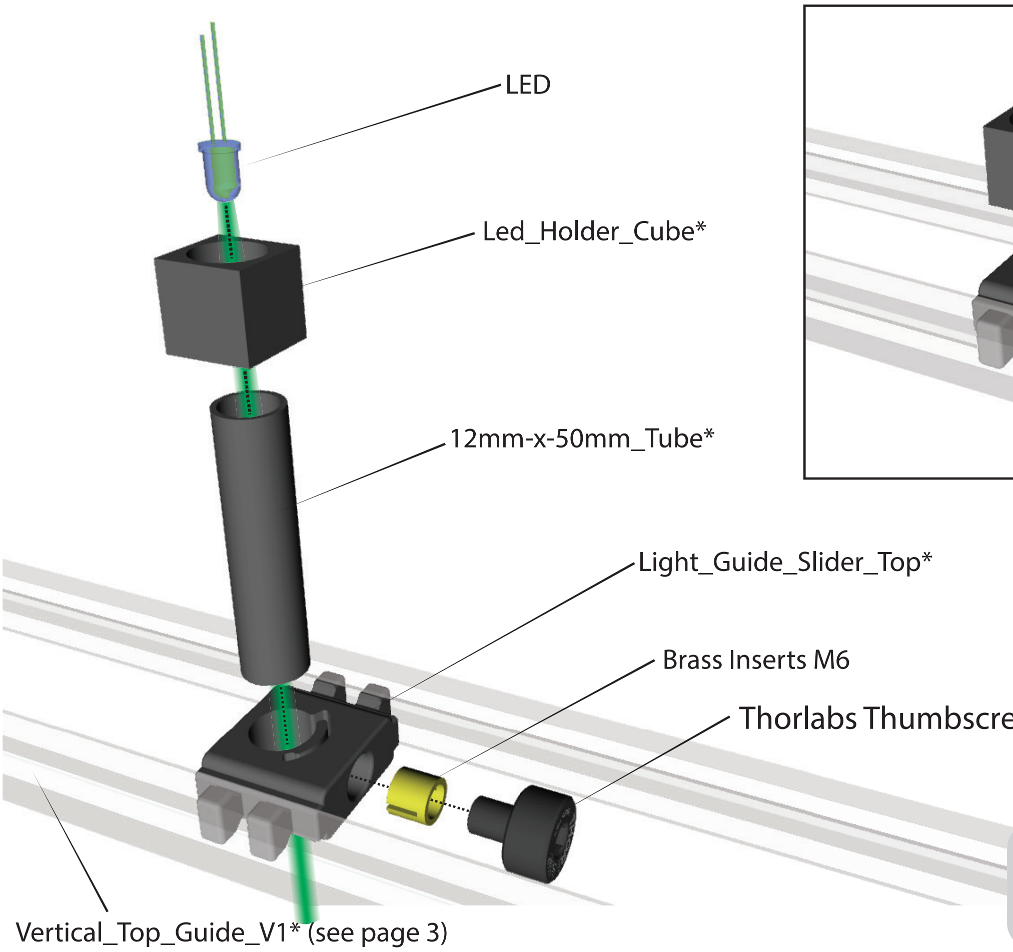

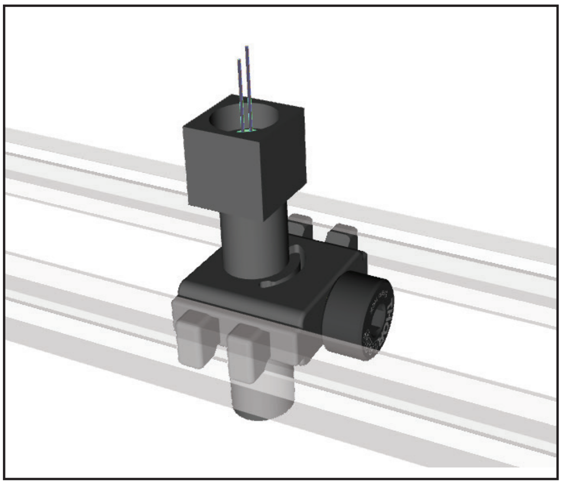

* 3D print with SLS technology

#### Neutral Density Filter Set

**Supplier: Reichmann Feinoptik**

| Density | Density tolerance | Transmission | Wavelength |
|---|---|---|---|
| 0.15 | +/-0.005 | 71.000% | 546nm |
| 0.30 | +/-0.005 | 50.120% | 546nm |
| 0.60 | +/-0.015 | 25.120% | 546nm |
| 1.00 | +/-0.03 | 10.000% | 546nm |
| 1.30 | +/-0.03 | 05.012% | 546nm |
| 1.60 | +/-0.03 | 02.512% | 546nm |
| 2.00 | +/-0.05 | 01.000% | 546nm |
| 2.30 | +/-0.05 | 00.501% | 546nm |
| 2.60 | +/-0.06 | 00.251% | 546nm |
| 3.00 | +/-0.06 | 00.100% | 546nm |
| 3.30 | +/-0.08 | 00.050% | 546nm |
| 3.60 | +/-0.08 | 00.025% | 546nm |
| 4.00 | +/-0.08 | 00.010% | 546nm |
| 4.30 | +/-0.08 | 00.005% | 546nm |
| 4.60 | +/-0.15 | 00.003% | 546nm |
| 5.00 | +/-0.15 | 00.001% | 546nm |

**Irradiance of White LEDs* with neutral density filters**

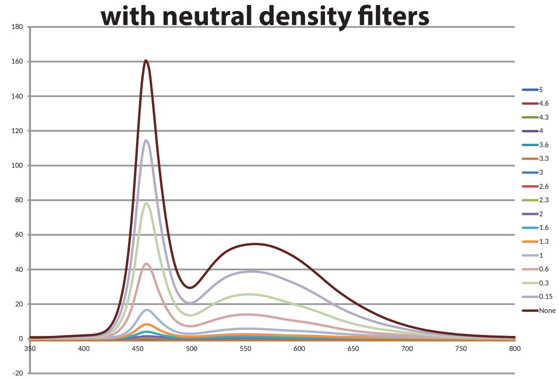

#### Omega Optical Dichroics, Beamsplitter and Mirror

##### Green Dichroic
565DRLPXR 201278 AOI 45˚

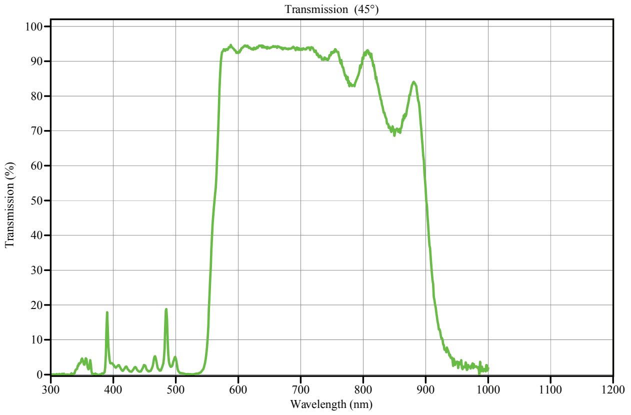

##### UV Dichroic
430DCLP 201276 AOI 45˚

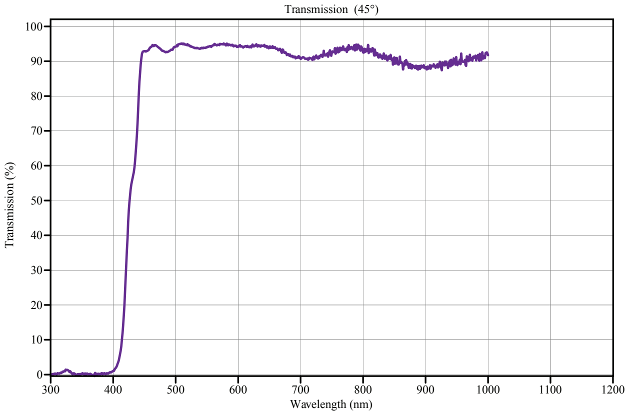

##### Blue Dichroic
505DRLP 201277 AOI 45˚

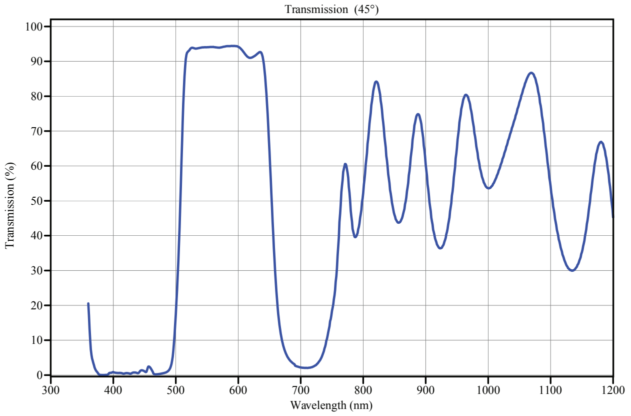

##### 50/50 Beamsplitter 201279 AOI 45˚

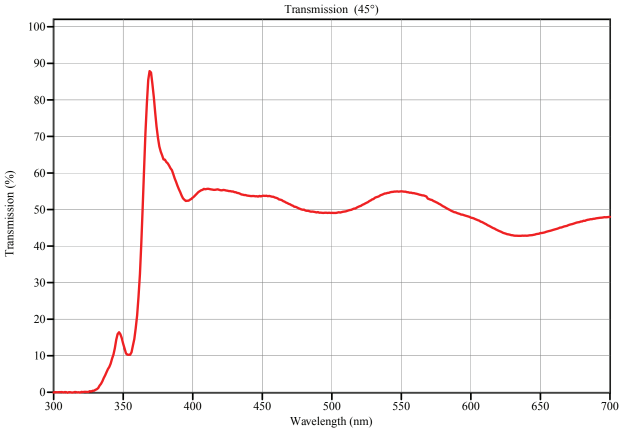

##### Mirror

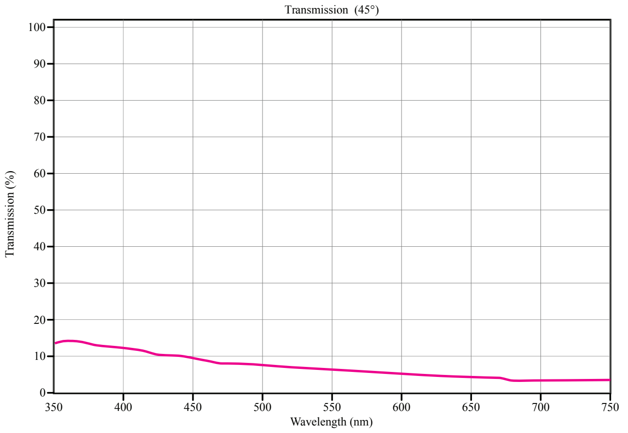
