## Supplementary material for "3D-printed Planktonic Observational Setup and Analysis Pipeline TrackmateTaxis": TrackmateTaxis Analysis Pipeline

### Step-by-Step Guide for TrackmateTaxis Software Pipeline

Contact Information
If you need any additional info, do not hesitate to contact me.

ccdoering

/Clemens_Doering

#### Video Recordings

- All recordings should be **uncompressed** and in **avi-format**
- To allow proper calibration it is necessary to crop each file to the width of the container. The margins above and below the container are irrelevant for the calibration.
- Alternatively, if the exact horizontal dimension are known, these can also be used for calibration

#### TrackmateTaxis - Tracking

Overview

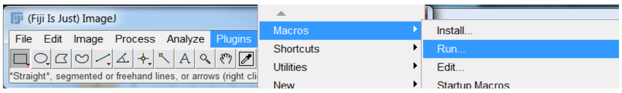


- This script runs in ImageJ and is written in Jython
- Upon script execution
- The first window asks for the working directory where the avi files are located
- The next window is the main user input interface:

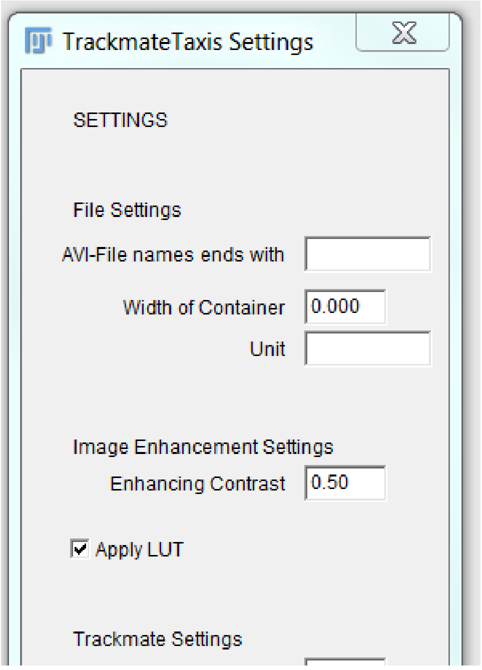


**Specify File ending:**

- If only a few files with the same ending in their filename are to be processed (*****cropped-nov.avi, ****cropped-jan.avi – filling in "nov" would only process one file) leave empty if not needed! Do not write ".avi", it would look for ".avi.avi"

**Width of container**

- If left empty, no calibration can be performed and all tracking will be done in pixels! The script will use the width of the recording in pixel and use the value for width of container to calibrate the recording.

**Unit**

- This is the unit used for the calibration

**Enhancing Contrast**

- Default: 0.5

**Apply LUT**

- Some videos have problems when applying LUT if it's a problem turn this off

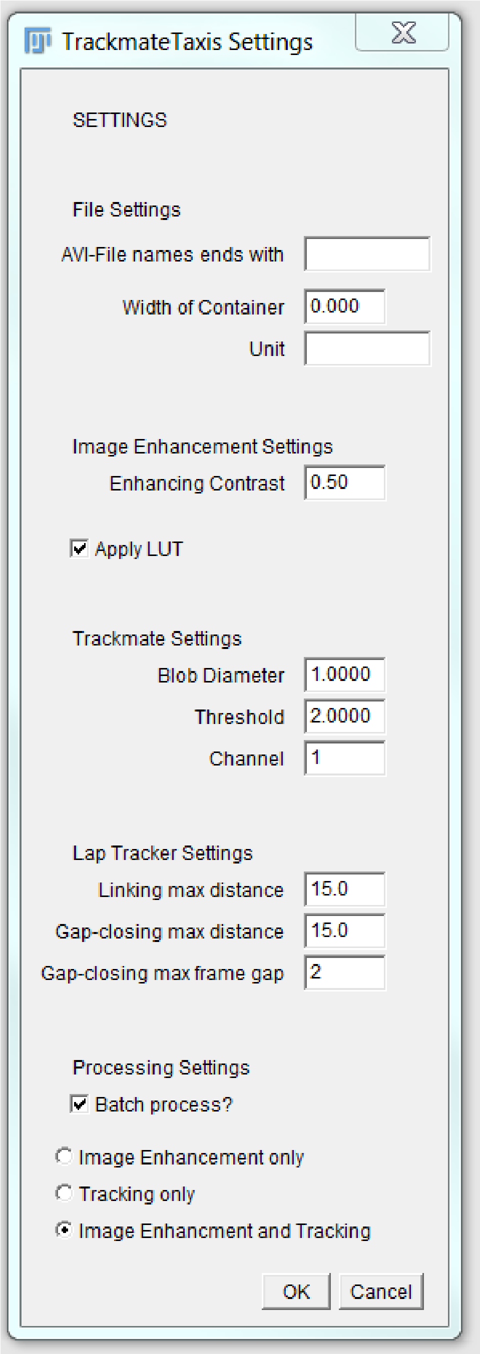


**Blob Diameter**

- This is the diameter of the particle to be tracked

**Threshold**

- Exclude objects otherwise detected

**Channel**

- If multiple channels are present

**Linking max distance**

- Maximum distance between any two points in different frames to be of one identity

**Gap-closing max distance**

- Maximum gap between any two points in different frames to be of one identity

**Gap-closing max frame gap**

- Maximum amount of frames allowed between two points to be considered of same identity

**Batch Process?**

- If only one file of all the files in the folder is to be processed, uncheck this box. Depending on the following settings this will alter the behavior of the script

**Image Enhancement only**

- This option will perform the image enhancement.
- If batch processing is chosen, the user is asked for the framerate of the videos and then each processed video is saved automatically in the working directory with the ending "EHN.avi"
- If batch processing is turned OFF it will process one video and show the result to the user. This is intended to be used to find the settings for the trackmate! Go to Plugins > Tracking > Trackmate and find good Blob Diameter and Threshold settings!

**Tracking only**

- Will use trackmate with the settings put into the gui without performing image enhancement

**Image Enhancement and Tracking**

- This will perform both actions

#### Finding the right settings/parameters

- The settings for tracking can be determined by running Image Enhancement Only and Batch mode turned off! After the image enhancement is complete, run Trackmate from the plugins in ImageJ and test the detection radius and threshold.
- Once settings are found rerun the script with the determined settings.
- The script will write a Logfile, named with the current Date and Time in the working directory. This will save all the settings put into the script and which files have been processed.
- The script output are two comma separated value (csv) files per video. All starting with the same date and time in their filename as the Logfile. One ending with trackstats.csv containing all information about the tracks with each of their positional information as well as some quality values. The second csv file ends with velocities.csv and contains all the velocity values of a particle has been tracked in consequent frames.

### Manual Tracking

It is possible to use Trackmate manually. After the tracking is completed, it is important to save the statistic files with the proper file name ending. The track statistics should be name *trackstats.csv and the link statistics as velocities.csv. The spot statistics is not needed for the subsequent part of the pipeline.

### Image Enhancement

The image enhancement performs the following steps:

- Conversion to 8 bit
- Enhancing contrast by value specified by user (default 0,5)
- Sharpen
- Apply LUT if chosen
- Invert
- Z-Projection (Average method)
- Subtraction of Z-Projection from each frame of the stack

### TrackmateTaxis - Analysis

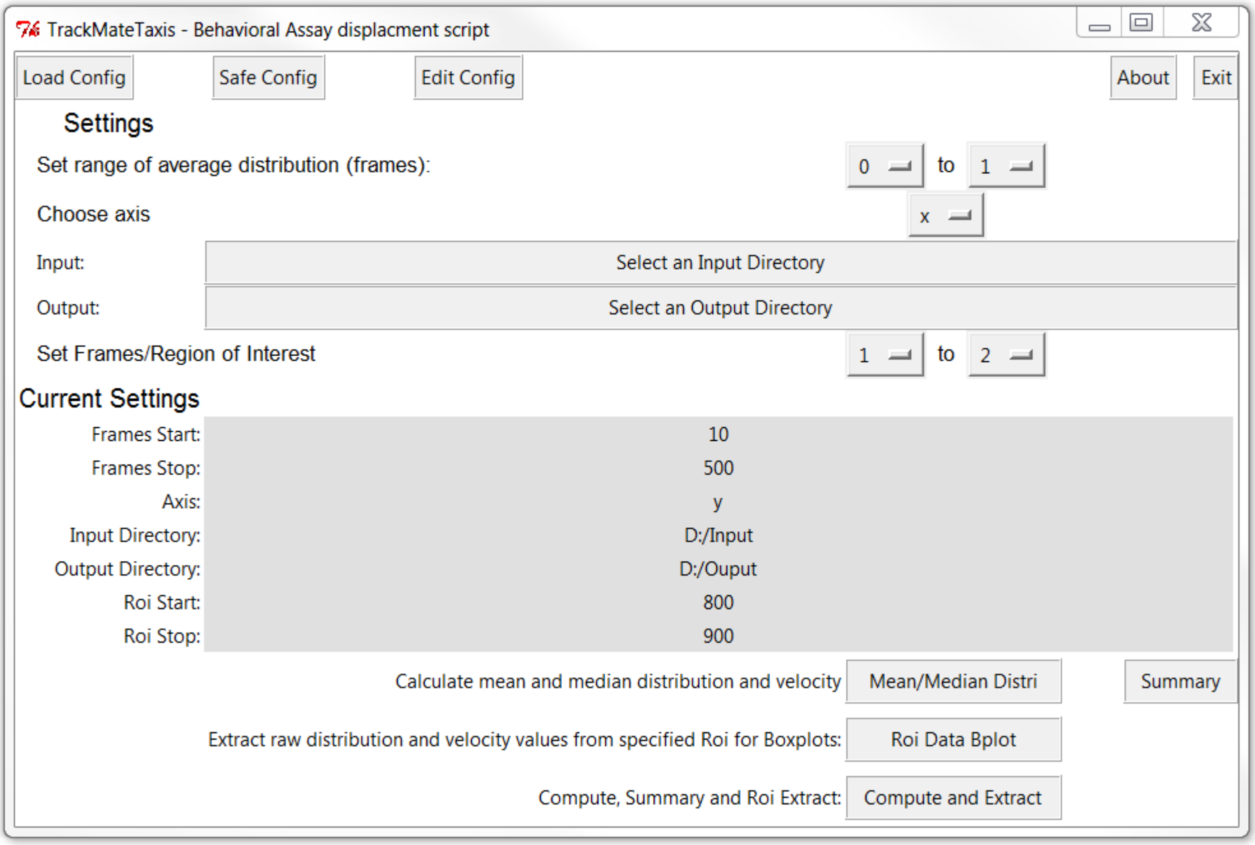

### Overview

- This script is a python script and needs a number of packages installed to run!
- We recommend to install Anaconda2 4.0.0 to ensure all required packages are installed. Newer version of Anaconda2 might remove needed packages!
- When the script is executed, it will open a Graphical User Interface (GUI).

**The GUI contains four sections.**

- The **header** is responsible for loading, saving and editing configuration files as well as closing the application. There is also an option to open a small "about" file for immediate help.
- The **input** section allows the user to choose input and output folder as well as set some parameters
- The **output** section shows the current settings the program will run*
- *if a configuration file has been edited and saved manually without restarting the software it will not update BUT will still work with the settings saved in the configuration file.
- The **footer section** allows the user to choose either to:
  - Calculate the mean and median distribution and velocity over the entire video. Will also plot the mean and median distribution.
  - Create a Summary of all files regarding distribution and velocity in the current OUTPUT directory.
  - Pool all values of the specified ROI into a single file and create a boxplot for all processed files for distribution and velocity. It will subtract the median value of the distribution as specified in the configuration file.
- The configuration file is where the script gets all its variables. If you need values that are not available through the gui, change them in the config file directly, it's just a text file!

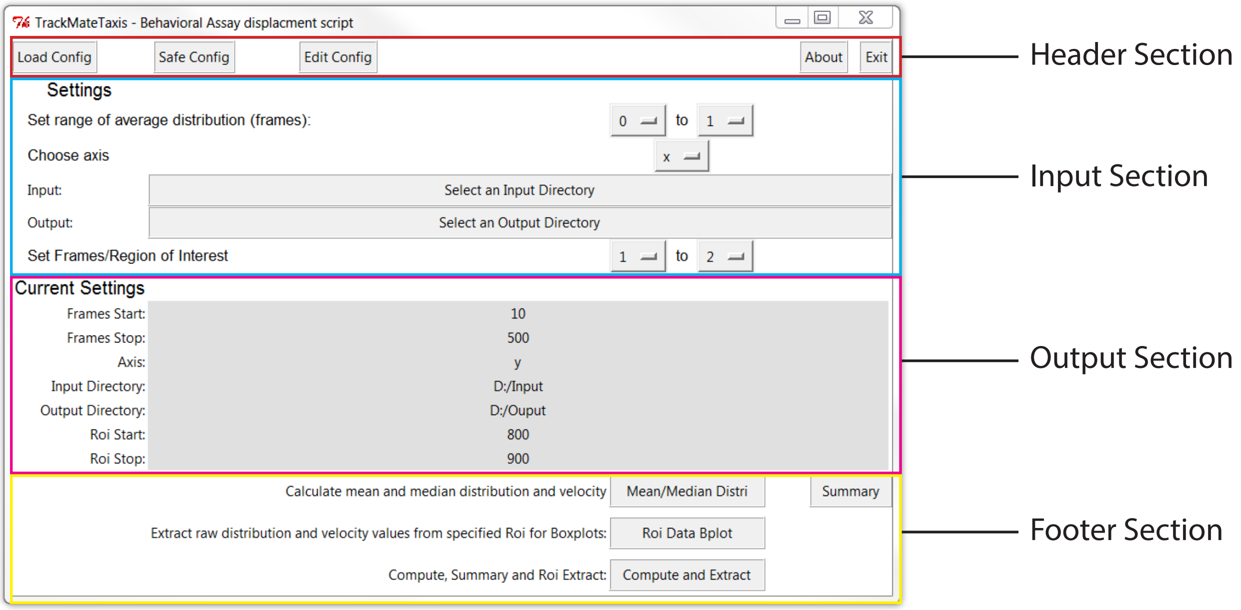

#### Parameters of the Input Section

##### Frames of Average Distribution

If no stimulus is present, it can be assumed that the animals are equally spread out in their container. Depending on swimming speed, animal density or other reason the very first frame might not be exactly evenly distributed. For this purpose it is possible to define a time frame where no stimulus is present and an even distribution can be determined by the mean/median position during this time frame.

##### Axis

The script will only look at one axis, x or y. This depends on the source of the stimulus or the major movements of the animals in response a stimulus.

##### Input Folder

All the files ending with *trackstats.csv and *velocities.csv need to be in here.

##### Output Folder

All the newly created excel files and pdfs with the plots will be saved here

##### Frames/Region of Interest

To obtain data to create boxplots it is important to get all the positional or velocity data for any given frame. These Frames of Interest or Region of Interest can be specified here. The script would then extract all the values that lie within range of the region of interest.

#### Configuration File

A configuration file is a simple text file that contains all the parameters the script should use. The script will write a default configuration file called "trackmatetaxis_config.txt" on first time initiation. It is possible to save the current configuration settings into a new file. It is also possible to Load different configuration files or open the current configuration file in use. If a configuration file is edited and saved manually it will not automatically update in the output view! Executing the script will still work with the manually saved settings! If someone needs to see the new settings either reload the configuration file by clicking on "Load Config " or restart the software to reload the default configuration file.

#### Footer Section

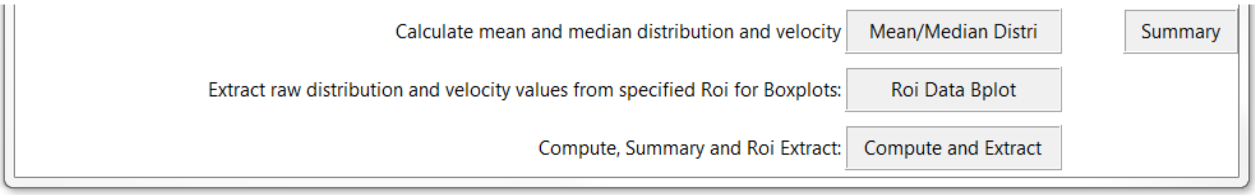


##### Mean/Median Distri

This part of the script will calculate the mean, quantiles (2%, 9%, 25%, 50%, 75%, 91% and 98%) and minimum and maximum positional changes as well as velocity for each frame. While the velocity is calculated by pooling all relevant values for each frame together, the positional changes are the difference between the mean/median position during the defined frames of average distribution and the mean/median of all the positional data for each frame. These adjusted values for the mean and quantiles are stored separately. The output for each video are two graphs one for mean and another for median distribution (including the 25% and 75% quantiles) as pdfs and two excel files for distribution and velocities each.

##### Summary

Summary will look in the current OUTPUT folder and look for filenames that are generated by the Mean/Median Distri function. It will matter which axis is selected in the current configuration file. It will extract all the adjusted values for each found excel file into a single excel file.

**Roi Data Bplot**
This part of the script will extract all positional data and velocity data for the selected range of frames (Region of Interest) and adjust the positional data in the same way that the Mean/Median Distri does by using the median of the frames of average distribution. It will look for the original tracking files in the INPUT folder. After all the values have been extracted from each input file it will create a summary file including all the files in the output folder that end with the same ROI settings as are currently set in the software. Finally, it will create a Boxplot with from all the combined files and save it as a pdf.

**Compute and Extract**
This function will execute Mean/Median Distri then Summary and finally the Roi Dat Blot one after the other.

#### SpectraRad Photon Counter Conversion

A few words on the script to convert from Irradiance to photon count. This script needs the output of spectrometer SpectraRad from B&Wtek. It will not work for any other unless they use the exact same csv output file!
If you want to use the script, open it in any text editing software and specify the LED range and working folder. Once completed, save the changes and execute the script in your python enviroment.
